# Intestinal fructose metabolism triggers a GLP-1–β-cell axis to prevent post-fructose hyperglycaemia

**DOI:** 10.1101/2025.04.01.646509

**Authors:** Naoya Murao, Yusuke Seino, Risa Morikawa, Shihomi Hidaka, Takuya Haraguchi, Eisuke Tomatsu, Mutsumi Habara, Tamio Ohno, Norihide Yokoi, Norio Harada, Yoshitaka Hayashi, Yuichiro Yamada, Atsushi Suzuki

**Author notes:** Corresponding author: Naoya Murao, M.D., Ph.D., Main address, Departments of Endocrinology, Diabetes and Metabolism, Fujita Health University, School of Medicine, 1-98 Dengakugakubo, Toyoake, Aichi 4701192 Japan. Institute for Diabetes and Organoid Technology (IDOT), Helmholtz Diabetes Center, Helmholtz Zentrum München Deutsches Forschungszentrum für Gesundheit und Umwelt (GmbH), Ingolstädter Landstraße 1, D-85764 Neuherberg, Germany. **PREPRINT PUBLICATION** This manuscript was first published as a preprint: Murao, N., Seino, Y., Morikawa, R., Hidaka, S., Haraguchi, T., Tomatsu, E., Habara, M., Ohno, T., Yokoi, N., Harada, N., Hayashi, Y., Yamada, Y., and Suzuki, A. (2025). Intestinal fructose metabolism/GLP-1/β-cell axis counteracts hyperglycemia after short-term fructose ingestion. bioRxiv.04.01.646509; doi: https://doi.org/10.1101/2025.04.01.646509.

## Abstract

Fructose ingestion increases circulating GLP-1 and insulin, yet the specific contributions of these hormonal responses to glycaemic control remain incompletely defined. We hypothesised that fructose metabolism in intestinal L-cells triggers GLP-1 secretion, which then potentiates insulin secretion and counteracts fructose-induced hyperglycaemia. To test this hypothesis, we systematically characterised metabolic responses across multiple mouse strains after 24 h ad libitum fructose ingestion. In both lean (NSY.B6-*a*/*a*) and obese diabetic (NSY.B6-*A*^*y*^/*a*) mice, fructose elevated plasma insulin, glucagon-like peptide 1 (GLP-1), and glucose-dependent insulinotropic polypeptide (GIP). The insulin response was preserved in GIP receptor-deficient mice (*Gipr*^−/−^) but was abolished in proglucagon-deficient mice (*Gcg*^−/−^) by pharmacological GLP-1 receptor antagonism, indicating a requirement for GLP-1, but not GIP. Across strains, fructose-induced insulin response correlated with attenuation of post-fructose glycaemia, consistent with insulin being essential for suppressing fructose-induced hyperglycaemia. To explore the mechanism underlying fructose-induced GLP-1 secretion, we combined ATP-sensitive potassium channel–deficient mice (*Kcnj11*^−/−^), GLUTag L-cell line, and metabolic tracing of ^13^C-labelled fructose in freshly isolated intestinal crypts. These complementary approaches support a model in which fructolysis increases the ATP/ADP ratio in L-cells, closes K_ATP_ channels, and stimulates GLP-1 secretion. In obese diabetic mice, increased fructolytic flux and a higher ATP/ADP ratio were associated with elevated GLP-1 levels, further corroborating this model. Collectively, our findings indicate that intestinal fructose metabolism drives GLP-1 secretion required to potentiate insulin secretion, thereby establishing a gut–pancreas axis that counter-regulates fructose-induced hyperglycaemia.

**KEY POINTS SUMMARY:**

- Fructose ingestion acutely increases plasma insulin levels, but the underlying mechanisms and physiological significance remain elusive.
- Our study demonstrates that short-term (24h) fructose ingestion in mice elevates both insulin and glucagon-like peptide 1 (GLP-1) levels in the blood, with the plasma insulin response being GLP-1-dependent.
- We found that fructose metabolism in intestinal L-cells triggered GLP-1 secretion by increasing the ATP/ADP ratio and closing ATP-sensitive K^+^ channels (K_ATP_ channels).
- This intestinal fructose metabolism/GLP-1/β-cell axis plays a crucial role in preventing fructose-induced hyperglycaemia, an effect that is compromised in obese diabetic mice.
- These insights highlight the previously unclear metabolic responses following short-term fructose ingestion and their importance in glucose homeostasis.

**ABSTRACT FIGURE LEGEND:** This study investigated the hormonal effects of short-term fructose consumption in mice, allowing them ad-lib access to fructose solution for 24 h. Fructose metabolism in intestinal L-cells increases the intracellular ATP/ADP ratio, leading to GLP-1 secretion via K_ATP_ channel closure and Ca^2+^ influx. GLP-1 promotes insulin secretion from pancreatic β-cells. The fructose metabolism/GLP-1/insulin pathway is essential for mitigation of fructose-induced hyperglycaemia. Figures were drawn using BioRender.com.

## 3. INTRODUCTION

Fructose is the second most abundant dietary monosaccharide after glucose (Larke et al., 2023). Multiple lines of evidence indicate that high intake of certain fructose-containing foods impairs metabolic health and contributes to type 2 diabetes (Stanhope et al., 2009; Imamura et al., 2015; Choo et al., 2018). Therefore, understanding the metabolic fate of fructose and the hormonal responses it elicits is critical. However, these processes remain less well-characterised than those for glucose.

In the small intestine, fructose enters enterocytes primarily through glucose transporter 5 (GLUT5), whereas glucose is absorbed predominantly through sodium–glucose cotransporter 1 (SGLT1), with a smaller contribution from apical GLUT2 under certain conditions. Both sugars are then exported across the basolateral membrane into the portal circulation by GLUT2 (Pepin et al., 2019). Blood glucose levels are tightly maintained within the normal physiological range owing to glucose-induced insulin secretion (GIIS) from pancreatic β-cells. GIIS is regulated by multiple mechanisms. Primarily, pancreatic β-cells directly sense blood glucose and secrete insulin through a series of events: glucose metabolism leading to an increase in the intracellular ATP/ADP ratio, closure of ATP-sensitive potassium channels (K_ATP_ channels), depolarisation of the cell membrane, opening of voltage-dependent calcium channels, and influx of Ca^2+^ that stimulates the exocytosis of insulin-containing granules (Henquin, 2000; Henuin, 2009). Additionally, gastrointestinal factors, most notably glucagon-like peptide 1 (GLP-1) and glucose-dependent insulinotropic polypeptide (GIP), further potentiate glucose-induced insulin secretion (GIIS). Following glucose ingestion, these incretins are secreted from L- and K-cells in the gut and activate their corresponding receptors in pancreatic β-cells (Seino et al., 2016; Santos-Hernández et al., 2024; Holst and Gromada, 2004). Mice lacking both incretin receptors have shown significant impairments in plasma insulin response and glucose tolerance following oral glucose administration (Hansotia et al., 2014; Preitner et al., 2004; Ahrén et al., 2021). Collectively, the incretin/β-cell axis is critical for glucose homeostasis (Nauck et al., 1986; Nauck and Meier, 2016).

Similar to glucose, fructose increases plasma GLP-1 and insulin levels in rodents (Kuhre et al., 2014; Seino et al., 2015) and healthy humans (Kong et al., 1999; Kuhre et al., 2014). However, the physiological significance of these responses remains poorly defined. We hypothesised that an incretin/β-cell axis analogous to that engaged by glucose also operates for fructose and helps maintain normoglycaemia after fructose ingestion. We tested this by characterising metabolic responses after short-term (24 h) ad libitum fructose intake across multiple mouse strains. We found that fructose metabolism in intestinal L-cells drives GLP-1 secretion, which is required to potentiate insulin secretion from β-cells. Together, we identified an intestinal pathway linking fructose metabolism, GLP-1, and β-cells that acts as a counterregulatory mechanism limiting fructose-induced hyperglycaemia.

## 4. METHODS

### 4.1. Mice

NSY.B6-*Tyr*+ *A*^*y*^/Hos mice, recently established by Ohno et al. (Ohno et al., 2022), were obtained from Hoshino Laboratory Animals (Ibaraki, Japan) and used for experiments at 19– 20 weeks of age. C57BL/6JJcl (RRID:IMSR_JCL:JCL:MIN-0003) was purchased from CLEA Japan (Tokyo, Japan) and employed for experiments at 13 weeks of age.

Colonies of *Gcg*^*gfp*/+^ and *Gcg*^*gfp*/*gfp*^ (hereafter referred to as *Gcg*^−/−^) mice (Hayashi et al., 2009), *Gipr*^−/−^ mice (Miyawaki et al., 1999), and *Kcnj11*^−/−^ mice (Miki et al., 1998) were maintained by intercrossing heterozygotes and homozygotes at Fujita Health University. All mice were housed under specific-pathogen-free conditions at 23 ± 2 °C and 55 ± 10% relative humidity with 12-hour light-dark cycles (8am–8pm), with ad libitum access to water and standard chow CE-2 (CLEA Japan, Tokyo, Japan). The health status of the mice was monitored regularly. All experiments were conducted using male mice. Body weight and blood glucose levels were measured at 9 am under ad libitum conditions. All in vivo experiments were performed with the approval of APU22080 by the Institutional Animal Care and Use Committee of Fujita Health University, in compliance with the Guidelines for Animal Experimentation at Fujita Health University and current Japanese legislation (Kagiyama et al., 2006). Efforts were made to minimise the stress and pain during the experiments.

Prior to the isolation of pancreatic islets (Section 4.9.) and intestine (Sections 4.16. and 4.18.), mice were euthanised by exposure to 5% isoflurane. Briefly, mice were placed in a sealed induction chamber (Shinano Seisakusho, Tokyo, Japan, Cat# SN-487-85) connected to an inhalation anaesthesia system (Shinano Seisakusho, Tokyo, Japan, Cat# SN-487). Isoflurane (FUJIFILM Wako Pure Chemical, Osaka, Japan, CAS: 26675-46-7, Cat# 095-06573) was vaporised and continuously delivered at a constant 5% concentration. Death was confirmed by the absence of spontaneous respiration and heartbeat for at least 2 min and was assured by bilateral thoracotomy and cardiac exsanguination prior to tissue collection.

### 4.2. Fructose administration

The schedule of fructose administration is shown in Fig. 1A. Mice were provided ad libitum access to 20% fructose as their sole water source for 24 h from 12:00 to 12:00. The control group received water. From 8am to 12am on the second day, the mice were deprived of chow and access to 20% fructose or water was maintained. The mice were subjected to fasting conditions with continuous access to fructose. Blood sampling and metabolic tests were conducted immediately after the administration period. For diazoxide treatment, diazoxide (Tokyo Chemical Industry, Tokyo, Japan, Cat# D5402) was pre-dissolved in dimethyl sulfoxide (DMSO) (FUJIFILM Wako Pure Chemical, Osaka, Japan, Cat# 041-29351) at a concentration of 1M and suspended in water or fructose solution at a final concentration of 10 mM, which was administered as described previously. For exendin [9-39] treatment, exendin [9-39] (Peptide Institute, Osaka, Japan, CAS: 133514-43-9, Cat# 4526-v) was dissolved in saline at 10 nmol/mL and injected intraperitoneally at a dose of 100 nmol/kg body weight immediately following a 24-hour fructose administration.

**Figure 1.**
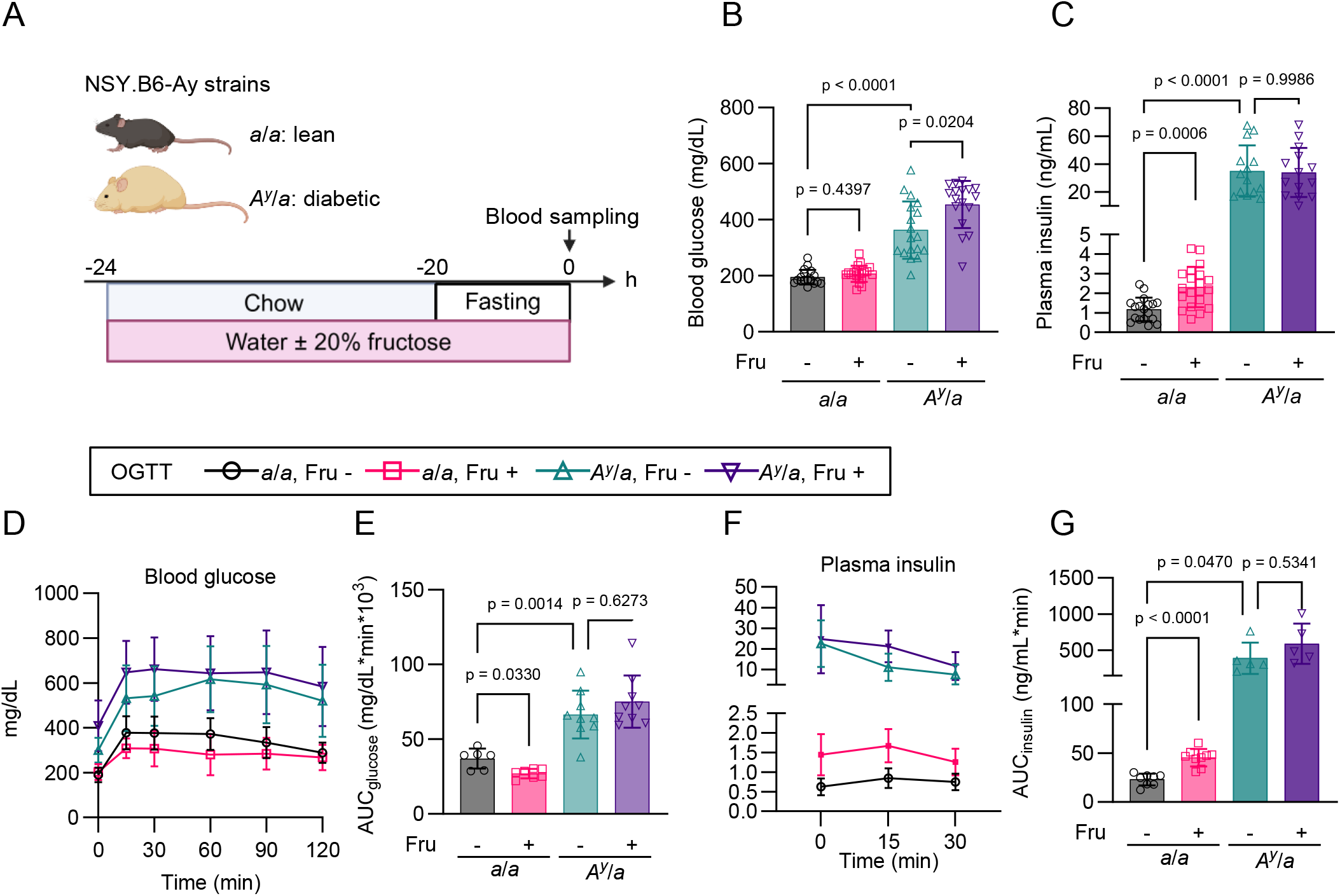
Short-term fructose ingestion in lean and diabetic mice. A. Schematic illustration of short-term fructose administration. Drawn using BioRender.com. B. Blood glucose levels following 24-hour fructose ingestion. *a*/*a* Fru −: n = 18; *a*/*a* Fru +: n = 20; *A*^*y*^/*a* Fru −: n = 18; *A*^*y*^/*a* Fru +: n = 17. N = 3. C. Plasma insulin levels following 24-hour fructose ingestion. *a*/*a* Fru −: n = 18; *a*/*a* Fru +: n = 20; *A*^*y*^/*a* Fru −: n = 14; *A*^*y*^/*a* Fru +: n = 14. N = 2. D. –G. Oral glucose tolerance test (OGTT) following 24-hour fructose ingestion. D, Blood glucose levels. E, Area under the curve (AUC) of blood glucose levels. *a*/*a* Fru-(circles): n = 6; *a*/*a* Fru + (squares): n = 6; *A*^*y*^/*a* Fru-(upward triangles): n = 6; *A*^*y*^/*a* Fru + (downward triangles): n = 6. N = 2. F, Plasma insulin levels. E, Area under the curve (AUC) of plasma insulin levels. *a*/*a* Fru −: n = 8; *a*/*a* Fru +: n = 9; *A*^*y*^/*a* Fru −: n = 5; *A*^*y*^/*a* Fru +: n = 5. N = 2. Data are presented as mean ± standard deviation (SD). Fru −: water-administered mice, Fru +: fructose-administered mice. Statistical comparisons were performed using Welch’s two-way ANOVA with Dunnett’s post-hoc test.

### 4.3. Measurement of plasma hormones

For hormone measurements, blood was collected from the tail vein of the unanesthetized mice using heparinised capillaries. Blood was supplemented with 4% BD-P800 (Becton, Dickinson and Company, Franklin Lakes, NJ, USA, Cat# 366421) dissolved in PBS and centrifuged at 2000 × g, 4 °C for 20 min to obtain plasma. Insulin, GIP, and glucagon levels were measured using Ultra-sensitive Mouse Insulin ELISA (Morinaga Bioscience, Yokohama, Japan, Cat# M1104), Rat/Mouse GIP (total) ELISA (Merck Millipore, Burlington, MA, USA, Cat# EZRMGIP-55K), and Glucagon ELISA Kit (Mercodia, Uppsala, Sweden, Cat# 10-1281-01), respectively. GLP-1 levels were measured using the V-PLEX GLP-1 (total) Kit (Meso Scale Discovery, Rockville, MD, USA, Cat# K1503PD) for Fig. A1E. Multi Species GLP-1 Total ELISA (Merck Millipore, Burlington, MA, USA, Cat# EZGLP1T-36K) was used for the other experiments. All ELISA were conducted in accordance with the manufacturer’s instructions.

### 4.4. Measurement of plasma fructose

To assess plasma fructose levels after 24-hour ingestion, blood was collected from the tail vein of unanesthetized mice, as described in Section 2.1. Alternatively, portal blood was collected immediately after the mice were euthanised by isoflurane exposure. To test acute fructose administration (Fig. A2, A–B), mice were fasted for 4 h with access to normal water. Aqueous fructose (2 g/10 mL) was administered via gavage at a dose of 2 g/kg body weight, and blood samples were obtained from the tail vein over 120 min. Blood was supplemented with 2 mM ethylenediaminetetraacetic acid (EDTA) and centrifuged at 2000 × g at 4 °C for 20 min to obtain plasma. For deproteinisation, 37 μL of plasma was diluted with 370 μL of acetonitrile, vortexed, and centrifuged at 15000 × g at 4 °C for 10 min. The supernatant was vacuum-dried and stored at −20 °C until analysis. Fructose was quantified using an LCMS-8060 Triple Quadrupole Liquid Chromatograph Mass Spectrometer (Shimadzu, Kyoto, Japan, RRID:SCR_020515). The sample diluent was formulated by mixing 10 mM aqueous ammonium acetate and acetonitrile at a ratio of 15:85. Samples were reconstituted with 20 μL sample diluent and spiked with 100 μM ^13^C_6_-fructose (Cambridge Isotope Laboratories, Tewksbury, MA, USA, CAS: 287100-63-4, Cat# CLM-1553-1) as an internal standard.

Separations were performed using a reverse-phase column Unison UK-Amino, 150 × 2 mm silica-gel column (Imtakt, Kyoto, Japan, Cat# UKA25) at 55 °C with the gradient method with water and acetonitrile as the mobile phase at a flow rate of 0.4 mL/min. Mass spectrometry was performed in the multiple reaction monitoring (MRM) negative mode for fructose (m/z 179.0 > 89.0) and ^13^C_6_-fructose (m/z 185.0 > 95.0).

### 4.5. Glucose and insulin tolerance tests

Glucose and insulin tolerance tests were performed as previously described (Murao et al., 2022). Following a 4-hour fast, aqueous glucose (1.5 g/10 mL) was administered via gavage at a dose of 1.5 g/kg body weight. Alternatively, human insulin (Humulin R, Eli Lilly, Indianapolis, IN, USA) was diluted with saline and administered intraperitoneally at a dose of 0.75 unit/kg body weight. Blood samples were collected from the tail vein over a 120-min period, during which mice were not allowed access to food and water. Blood glucose levels were measured using an Antsense Duo glucose analyzer (Horiba, Kyoto, Japan, Cat# LP-150).

### 4.6. Cell lines

GLUTag cells (Drucker et al., 1994) were acquired from Dr. Akira Hirasawa (Kyoto University), with permission from Professor Daniel Drucker. GLUTag cells were cultured in Dulbecco’s modified Eagle’s medium (DMEM) containing 1000 mg/L glucose (Sigma-Aldrich, St. Louis, MO, USA, Cat# D6046) supplemented with 10% FBS (BioWest, Nuaillé, France, Cat# S1400-500). All cells were maintained in a humidified incubator at 37 °C with 5% CO2.

### 4.7. Plasmids

GW1-PercevalHR was a generous gift from Professor Gary Yellen (Addgene plasmid # 49082; http://n2t.net/addgene:49082; RRID: Addgene_49082) (Tantama et al., 2013).

### 4.8. Reagents for in vitro experiments

For all in vitro experiments, Krebs-Ringer bicarbonate buffer-HEPES (KRBH: 133.4 mM NaCl, 4.7 mM KCl, 1.2 mM KH2PO4, 1.2 mM MgSO4, 2.5 mM CaCl2, 5 mM NaHCO3, 10 mM HEPES) supplemented with 0.1% bovine-serum albumin (Sigma-Aldrich, St. Louis, MO, USA, Cat# A6003) adjusted to pH 7.4 was used. Glucose was not added to KRBH unless otherwise specified. D-Fructose (FUJIFILM Wako Pure Chemical, Osaka, Japan, CAS: 57-48-7, Cat# 127-02765), Human GLP-1 7-36 amide (Peptide Institute, Osaka, Japan, CAS: 107444-51-9, Cat# 4344-v), and Human GIP (Peptide Institute, Osaka, Japan, CAS: 100040-31-1, Cat# 4178-s) were added to the KRBH only during the stimulation period. Nifedipine (FUJIFILM Wako Pure Chemical, Osaka, Japan, Cat# 14505781) and TEPP-46 (Selleck Chemicals, Houston, TX, USA, CAS: 1221186-53-3, Cat# S7302) were added during the pre-incubation and stimulation periods. GLP-1 and GIP were stored as a 100 μM solution in 0.1% aqueous acetic acid. Small-molecule reagents were stored as 1000× concentrate in dimethyl sulfoxide (DMSO) (FUJIFILM Wako Pure Chemical, Osaka, Japan, Cat# 041-29351). The reagents were diluted with KRBH immediately before the experiment. An equivalent volume of DMSO was added to the control when appropriate.

### 4.9. Isolation of pancreatic islets from mice

Digesting solution was formulated by supplementing 0.1w/v% Collagenase from Clostridium histolyticum (Sigma-Aldrich, St. Louis, MO, USA, Cat# C6885) to Hanks’ balanced salt solution (136.9 mM NaCl, 5.4 mM KCl, 0.8 mM MgSO_4_, 0.3 mM Na_2_HPO_4_, 0.4 mM KH_2_PO_4_, 4.2 mM NaHCO_3_, 10 mM HEPES, 1.3 mM CaCl_2_, 2 mM glucose). The mice were euthanised by exposure to 5% isoflurane. The pancreas was digested by 10-min incubation at 37 °C following intraductal injection of digesting solution. Islets were hand-picked and separated from exocrine tissues, transferred to 60-mm non-treated plates (AGC Techno Glass, Shizuoka, Japan, Cat# 1010-060), and cultured overnight in RPMI-1640 (Sigma-Aldrich, St. Louis, MO, USA, Cat# R8758) supplemented with 10% FBS (BioWest, Nuaillé, France, Cat# S1400-500) and 1% penicillin-streptomycin solution (FUJIFILM Wako Pure Chemical, Osaka, Japan, Cat# 168-23191) at 37 °C with 5% CO_2_ before the experiments.

### 4.10. Insulin secretion from islets

Insulin secretion from isolated islets was measured using the static incubation method, as previously described (Murao et al., 2022; Murao et al., 2025). Specifically, five size-matched islets were placed in each well of a 96-well plate (Corning, Glendale, AZ, USA, Cat# 353072) and were pre-incubated with 50 μL/well of KRBH supplemented with 2.8 mM glucose. Subsequently, 50 μL of KRBH supplemented with 19.4 mM glucose and 2 nM GLP-1 or GIP were added to the reaction, and the islets were incubated for an additional 30 min. The supernatant was subjected to insulin quantification using a homogeneous time-resolved fluorescence assay (HTRF) Insulin Ultrasensitive kit (Revvity, Waltham, MA, USA, Cat# 62IN2PEH) in accordance with the manufacturer’s protocol.

### 4.11. Transfection small interfering RNAs (siRNAs) targeting *Khk* and *Aldob*

siRNAs targeting *Khk* (Dharmacon, Lafayette, CO, USA, Cat# M-062217-01-0005), *Aldob* (Cat# M-064045-00-0005), and non-targeting siRNA (Cat# D-001206-14-50) were reverse-transfected using the DharmaFECT 2 transfection reagent (Dharmacon, Lafayette, CO, USA, Cat# T-2002-03). A complex of siRNA (40 nM) and DharmaFECT 2 (0.4%) was prepared in serum-free DMEM (Sigma-Aldrich, St. Louis, MO, USA, Cat# D5796) in accordance with the manufacturer’s instructions. For GLP-1 secretion measurements, siRNA/DharmaFECT 2 complex (100 μL/well) was combined with the cell suspension (0.75 × 10^6^ cells/mL, 400 μL/well) and seeded in a 24-well plate pre-coated with 3% Matrigel (Corning, Glendale, AZ, USA, Cat# 356234). For RT-PCR, siRNA/DharmaFECT 2 complex (200 μL/well) was combined with the cell suspension (0.75 × 10^6^ cells/mL, 800 μL/well) and seeded in a 12-well plate pre-coated with 3% Matrigel. For ATP/ADP ratio and Ca^2+^ measurements, the cells were seeded in a 35 mm glass-bottom dish (Matsunami Glass, Osaka, Japan, Cat# D11530H) pre-coated with 3% Matrigel at a density of 1.28 × 10^5^ cells/dish. Following 1 h incubation, the culture media was replaced with a mixture of siRNA/DharmaFECT 2 complex (400 μL/dish) and complete culture media (1.6 mL/dish). The subsequent experiments were conducted after a 48-hour culture period.

### 4.12. GLP-1 secretion from GLUTag cells

GLP-1 secretion from GLUTag cells was measured using the static incubation method. A 24-well plate (Corning, Glendale, AZ, USA, Cat# 353047) was pre-coated with 3% Matrigel (Corning, Glendale, AZ, USA, Cat# 356234). Cells were seeded at a density of 3 × 10^5^ cells/well and cultured for 48 h. KRBH was supplemented with 10 μM sitagliptin phosphate (Tokyo Chemical Industry, Tokyo, Japan; CAS: 654671-78-0, Cat# I1231) to inhibit the degradation of active GLP-1. The cells were washed three times with KRBH, followed by a pre-incubation period of 30 min with 300 μL/well KRBH. Subsequently, the supernatant was replaced with 300 μL/well of fresh KRBH containing the specified stimulations and incubated for 60 min at 37 °C. The reaction was terminated by cooling the plate on ice for ten minutes, after which the entire supernatant was collected for the quantification of released GLP-1 using the homogeneous time-resolved fluorescence assay (HTRF) active GLP-1 detection kit (Revvity, Waltham, MA, USA, Cat# 62 62GLPPEG) in accordance with the manufacturer’s instructions. Fluorescence was measured using an Infinite F Nano+ microplate reader (Tecan, Zürich, Switzerland).

### 4.13. Imaging of intracellular Ca^2+^ in GLUTag cells

Imaging of intracellular Ca^2+^ was performed as previously described (Murao et al., 2024), with minor modifications. The cells were seeded in a 35 mm glass-bottom dish (Matsunami Glass, Osaka, Japan, Cat# D11530H) pre-coated with 3% Matrigel (Corning, Glendale, AZ, USA,Cat# 356234) at a density of 1.28 × 10^5^ cells/dish and cultured for 48 h. Subsequently, the cells were loaded with 1 μM Fluo-4 AM (Dojindo, Kumamoto, Japan, Cat# F312) in KRBH supplemented with 0.04% Pluronic F-127 (Sigma-Aldrich, St. Louis, MO, USA, CAS: 9003-11-6, Cat# P2443-250G) for 20 min at 37 °C. Following brief washing, cells were loaded with 1 mL of KRBH, and basal recordings were performed for 300 s (from time −300 to 0). Immediately after the addition of 1 mL of KRBH supplemented with 20 mM fructose, recordings were resumed for another 1800 s. Time-lapse images were obtained using a Zeiss LSM 980 Airyscan2 inverted confocal laser scanning super-resolution microscope equipped with a Plan Apo 40×, 1.4 Oil DICII objective lens (Carl Zeiss Microscopy, Jena, Germany). The cells were excited at 488 nm laser with 0.3% output power, and fluorescence emission was measured at 508-579 nm. During observation, the cells were maintained at 37 °C using an incubator XLmulti S2 DARK (Pecon, Erbach, Germany). Images were acquired in the frame mode at a rate of 2 frames per second and with an image size of 212.2 × 212.2 μm (512 × 512 pixels). The obtained images were analysed using the ZEN 3.0 imaging software (Carl Zeiss Microscopy, Jena, Germany, RRID:SCR_021725). Cells were randomly chosen for analysis for each stimulation, and the number of cells analysed is indicated in the figure legends. The fluorescence intensity of the entire cell body (F) was calculated and normalised to its average between −5 and 0 min (F0).

### 4.14. Imaging of intracellular ATP/ADP ratio in GLUTag cells

The cells were seeded in a 35 mm glass-bottom dish (Matsunami Glass, Osaka, Japan, Cat# D11530H) at a density of 1.28 × 10^5^ cells/dish and cultured for 1 h. Subsequently, the cells were transfected with GW1-PercevalHR plasmid (0.1 μg/dish) or double transfection of GW1-PercevalHR and siRNAs using DharmaFECT 2 transfection reagent (Dharmacon, Lafayette, CO, USA, Cat# T-2002-03). Following a 48h culture, the cells were washed and loaded with 1 mL of fresh KRBH, and basal recordings were conducted for 300 s (from time −300 to 0). Immediately after the addition of 1 mL of KRBH supplemented with stimulations at a 2× concentration, recordings were resumed for an additional 2400 s with a time interval of 5 s. The cells were excited at 445 nm and 488 nm with 0.8% and 0.3% output power, respectively, and fluorescence emission was measured at 508-543 nm. The ratio (R) of the fluorescence at 445 and 488 nm was calculated and normalised to the average fluorescence intensity between −300 and 0 s (R0). The remaining experimental configurations were identical to those described for Ca^2+^ imaging.

### 4.15. Quantitative reverse transcription polymerase chain reaction (RT-qPCR)

cDNA was prepared from GLUTag cells using CellAmp Direct Lysis and RT set (Takara Bio, Shiga, Japan, Cat# 3737S/A) according to the manufacturer’s instructions. Quantitative real-time PCR was performed on a QuantStudio 7 Flex system (Thermo Fisher Scientific, Waltham, MA, USA, RRID:SCR_020245) using TaqMan Universal Master Mix II with UNG (Thermo Fisher Scientific, Waltham, MA, USA, Cat# 4440038) and Taqman probes: *Khk* (Cat# Mm00434647_m1), *Aldob* (Cat# Mm00523293_m1), and *Tbp* (Cat# Mm01277042_m1). Relative gene expression was calculated using the 2^−ΔΔCT^ method and normalised to *Tbp*.

### 4.16. Isolation of intestinal crypts from mice

The mice were fed standard chow and water until euthanasia by exposure to 5% isoflurane. Intestinal crypts were isolated from the distal half of the small intestine, including the jejunum and ileum, where fructose is absorbed (Bode et al., 1981). Isolation was conducted in accordance with established protocols, with minor modifications (Bas et al., 2014; O’Rourke et al., 2016). The distal half of the small intestine was longitudinally incised, rinsed with ice-cold phosphate-buffered saline (PBS), and sectioned into 5 mm segments using razors. The following procedure was conducted on ice. The intestinal fragments from two mice were combined in a 50 mL tube and dissociated into crypts through the following procedures: washing by vigorous pipetting in 10 mL PBS supplemented with 5 mM ethylenediaminetetraacetic acid (EDTA); digestion by gentle agitation in 10 mL 5 mM EDTA/PBS for 10 min; gentle agitation in 10 mL fresh 5 mM EDTA/PBS for an additional 30 min; gentle washing with 10 mL PBS; dispersion of crypts by vigorous pipetting with 10 mL PBS three times, with the supernatant combined; centrifugation at 300 × g for 5 min; the crypts were obtained as suspension after careful removal of the supernatant. The crypts isolated from two mice per genotype were combined and immediately utilised for subsequent metabolic labelling procedures. Fluorescent images were acquired using a BZ-X810 microscope (Keyence, Osaka, Japan, RRID:SCR_025160).

### 4.17. Fructose metabolic tracing in intestinal crypts

500 μL of the crypt suspension was aliquoted into 1.5 mL screw tubes and subsequently washed three times with 2.8 mM glucose-KRBH. The crypts were pre-incubated with 500 μL 2.8 mM glucose-KRBH for 60 min at 37 °C. Subsequently, 500 μL of 2.8 mM glucose-KRBH with or without 100 mM ^13^C_6_-fructose (Cambridge Isotope Laboratories, Tewksbury, MA, USA, CAS: 287100-63-4, Cat# CLM-1553-1) was added to the reaction (^13^C_6_-fructose final concentration: 50 mM). The crypts were incubated for an additional 60 min at 37 °C. To facilitate ventilation, the tube lids were kept open during incubation. Following centrifugation at 500 × g for 1 min, the supernatant was discarded, the crypt pellet was rapidly lysed by the addition of 500 μL ice-cold extraction buffer (67.5% methanol, 7.5% chloroform, and 25% water), and the entire tube was snap-frozen in liquid nitrogen. For extraction of the metabolites, samples were supplemented with 80 μL of diluted (1:640) internal standard (Human Metabolome Technologies, Yamagata, Japan, Cat# H3304-1002), 165 μL of methanol, and 465 μL of chloroform. The samples were then homogenised using a pre-cooled bead crusher at 3200 rpm for 1 min and centrifuged at 15000 × g at 4 °C for 3 min. The aqueous layer was transferred to pre-wetted ultrafiltration tubes (Human Metabolome Technologies, Yamagata, Japan, Cat# UFC3LCCNB-HMT) and centrifuged at 9100 × g, 4 °C until completely filtered. The filtrate was freeze-dried, re-dissolved in 10 μL of water, and subjected to mass spectrometry. The organic layer was evaporated by decompression at room temperature, and the residue was resuspended in lysis buffer (see Section 4.7.), which was then subjected to a BCA protein assay (Thermo Fisher Scientific, Waltham, MA, USA, Cat# 23225).

The concentration of metabolites was measured using G7100A capillary electrophoresis (Agilent Technologies, Santa Clara, CA, USA) interfaced with a G6224A time-of-flight LC/MS mass spectrometer (Agilent Technologies, Santa Clara, CA, USA). A G1310A isocratic pump (Agilent Technologies, Santa Clara, CA, USA) equipped with a G1379B degasser (Agilent Technologies, Santa Clara, CA, USA) was used to supply sheath liquid (Human Metabolome Technologies, Yamagata, Japan, Cat# H3301-2020). The mass spectrometer was operated in the negative ionisation mode. All separations were performed on fused silica capillaries (Human Metabolome Technologies, Yamagata, Japan, Cat# H3305-2002) at 25 °C using an anion analysis buffer (Human Metabolome Technologies, Yamagata, Japan, Cat# H3302-2021) as the background electrolyte. The applied voltage was set to 30 kV at 20 °C, together with a pressure of 15 mbar. Sheath liquid was delivered to a nebulizer by an isocratic pump at 1 mL/min. Chromatograms and mass spectra were analysed using MassHunter qualitative analysis version 10.0 (Agilent Technologies, Santa Clara, CA, USA, RRID:SCR_015040). Annotation and quantification of chromatogram peaks were performed using a standard mixture (Human Metabolome Technologies, Yamagata, Japan, Cat# H3302-2021) and D-fructose 1-phosphate dipotassium salt (Santa Cruz Biotechnology, Dallas, TX, USA, CAS: 103213-47-4, Cat# sc-500907) as references.

### 4.18. Immunohistochemistry

Tissue staining was performed as previously described (Nishida et al., 2024). Briefly, the distal half of the small intestine was collected from mice as described in Section 4.16. The tissue was fixed with 4% paraformaldehyde (FUJIFILM Wako Pure Chemical Co., Osaka, Japan, Cat# 161-20141) for 7 days and then embedded in paraffin. Sections (4 μm) were taken and de-paraffinised using the conventional method. Antigen retrieval was performed by boiling the de-paraffinised sections in sodium citrate buffer (10 mM sodium citrate, 0.1% NP-40, pH 6.0) in a microwave oven for 15 min. The sections were blocked with PBS supplemented with 10% goat serum and 0.1% Triton-X. Primary antibodies and dilutions: monoclonal mouse Anti-KHK Santa Cruz Biotechnology, Dallas, TX, USA, Cat# sc-377411) 1:100; polyclonal rabbit anti-GLP-1 (Abcam, Cambridge, UK, Cat# ab22625, RRID: AB_447206) 1:500. The primary antibodies were diluted in PBS containing 3% BSA and 0.01% Triton-X. Pancreatic sections were loaded with diluted antibodies and incubated overnight at 4 °C. Secondary antibodies: goat anti-rabbit IgG (H+L) Alexa Fluor 488 (Thermo Fisher Scientific, Waltham, MA, USA, Cat# A-11008, RRID: AB_143165) and goat anti-mouse IgG (H+L) Alexa Fluor 594 (Thermo Fisher Scientific, Waltham, MA, USA, Cat# A-11032, RRID: AB_2534091). The secondary antibodies were diluted 1:500 in PBS containing 1% goat serum and 0.01% Triton-X. The sections were then loaded with diluted secondary antibodies and incubated in the dark for 1 h at room temperature, followed by treatment with DAPI (Dojindo) diluted with PBS (1:2000) in the dark for 3 min at room temperature. Slides were mounted in Fluoromount (Diagnostic BioSystems, Pleasanton, CA, USA, Cat# K024). Images were acquired using a BZ-X810 microscope (Keyence, Osaka, Japan, RRID:SCR_025160) with a 20X lens. Optical configurations, including light intensity and exposure time (KHK: 1/6 s; GLP-1:1/6 s; DAPI: 1/60 s), were equal for *a*/*a* and *A*^*y*^/*a* samples. Grayscale images were generated using ImageJ ver. 1.53k (National Institute of Health, https://imagej.nih.gov/ij/index.html, RRID:SCR_003070).

### 4.19. Statistical Analysis

Sample sizes were estimated from the expected effect size based on previous experiments. No randomization or blinding was employed. Experiments were conducted multiple times on separate occasions using distinct cell passages or different mice, with N indicating the number of repetitions. For in vivo experiments and immunohistochemistry, n represents the number of mice. For GLP-1 secretion, n represents the number of different wells under the same conditions. For insulin secretion from islets, n represents the number of wells, each containing five islets. For Ca^2+^ and ATP/ADP ratio measurements, n represents the number of different single cells analysed, with data pooled from 2 to 3 independent measurements. For fructose metabolic tracing, n represents the number of samples aliquoted from crypts pooled from two mice per genotype. Plotted values included all data pooled across these independent repetitions. Data are presented as the mean ± standard deviation (SD) along with the plot of individual data points. For statistical comparisons between two groups, a two-tailed unpaired Welch’s unpaired t-test was used. For comparisons between three groups, Welch’s one-way analysis of variance (ANOVA) was followed by pairwise comparisons corrected using Dunnett’s method. Normality of the distribution was confirmed by the Shapiro-Wilk test. P-values were indicated in the figures, with those less than 0.05 considered statistically significant. Statistical analyses are indicated in the figure legends. Statistical analyses were performed using GraphPad Prism 10 (Graphpad Software, Boston, MA, USA, https://www.graphpad.com; RRID:SCR_002798).

## 5. RESULTS

### 5.1. Short-term fructose ingestion increases plasma insulin levels in lean mice

To examine the metabolic impact of fructose in both lean and diabetic states, we used the NSY.B6-*A*^*y*^ mouse strain, a model of spontaneous obese diabetes. These strains were recently developed by introducing an obesity-associated agouti yellow (*A*^*y*^) mutation in the agouti (a) gene into a diabetes-susceptible NSY background (Ohno et al., 2022). Male mice with the heterozygous (*A*^*y*^/*a*) genotype exhibited obesity and elevated blood glucose levels, whereas their wild-type (*a*/*a*) counterparts remained lean (Fig. A1, A–B), which is consistent with previous findings (Ohno et al., 2022). Fructose was administered as illustrated in Fig. 1A.

Following the 24h administration period, *A*^*y*^/*a* mice exhibited significant hyperglycaemia and hyperinsulinaemia compared to *a*/*a* mice, both without fructose (Fig. 1, B–C). In *a*/*a* mice, fructose did not affect blood glucose (Fig. 1B) but increased plasma insulin levels (Fig. 1C). Conversely, in *A*^*y*^/*a* mice, fructose increased blood glucose (Fig. 1B) but did not alter plasma insulin levels (Fig. 1C).

During oral glucose tolerance tests (OGTT), *A*^*y*^/*a* mice exhibited impaired glucose tolerance (Fig. 1, D–E) and hyperinsulinemia compared to *a*/*a* mice, both without fructose (Fig. 1, F–G). In *a*/*a* mice, fructose ingestion resulted in improved glucose tolerance (Fig. 1, D–E), accompanied by elevated insulin levels persisting after glucose administration (Fig. 1, F–G). Conversely, fructose ingestion did not elicit any significant differences in glucose tolerance or insulin levels in *A*^*y*^/*a* mice (Fig. 1, D–G). In intraperitoneal insulin tolerance tests, *A*^*y*^/*a* mice exhibited severe insulin resistance, with fructose having no effect on either strain (Fig. A1, C–D). These results suggest that improved glucose tolerance is primarily attributable to enhanced insulin response.

These findings highlight the differential metabolic responses following fructose ingestion under lean and diabetic conditions: fructose-induced insulin response may help maintain normal glycaemia and enhance glucose tolerance in lean mice, whereas these effects were abolished in diabetic mice.

### 5.2. GLP-1 is required for fructose-induced insulin response

We investigated circulating fructose and fructose-stimulated insulinotropic hormone levels as potential factors stimulating fructose-induced insulin secretion in vivo.

Fructose enhances glucose-induced insulin secretion from β-cells in vitro at high concentrations (3–27 mM) (Grodsky et al. 1963; Zawalich et al. 1977; Grant et al. 1980; Kyriazis et al. 2012; Bartley et al. 2019; Murao et al. 2025). To assess whether plasma fructose levels attained these concentrations under our conditions, fructose in the tail and portal veins was quantified using liquid chromatography-tandem mass spectrometry (LC-MS/MS). Without fructose administration, fructose concentrations were approximately 5 μM in both veins. However, fructose ingestion elevated the concentrations to 100–200 μM and 10–20 μM in the tail and portal veins, respectively (Fig. 2, A–B). Fructose levels in the portal vein were lower and exhibited greater variability than those in the tail vein. This is likely due to the random timing of fructose consumption followed by its swift removal from the portal blood (Jang et al., 2018). We also tested the acute gavage of 2 g/kg fructose to *a*/*a, A*^*y*^/*a*, and C57BL/6J (B6) mice (Fig. A2, A–B). Plasma fructose levels reached their highest point (~100 μM) between 15 and 30 min, returning to initial levels within 120 min, indicating that the maximum plasma fructose level was approximately 100 μM. Nevertheless, our recent findings showed that fructose concentrations below 3 mM failed to affect GIIS in *a*/*a* mouse islets (Murao et al., 2025). The difference between these concentrations does not support the notion that circulating fructose directly stimulates β-cells.

**Figure 2.**
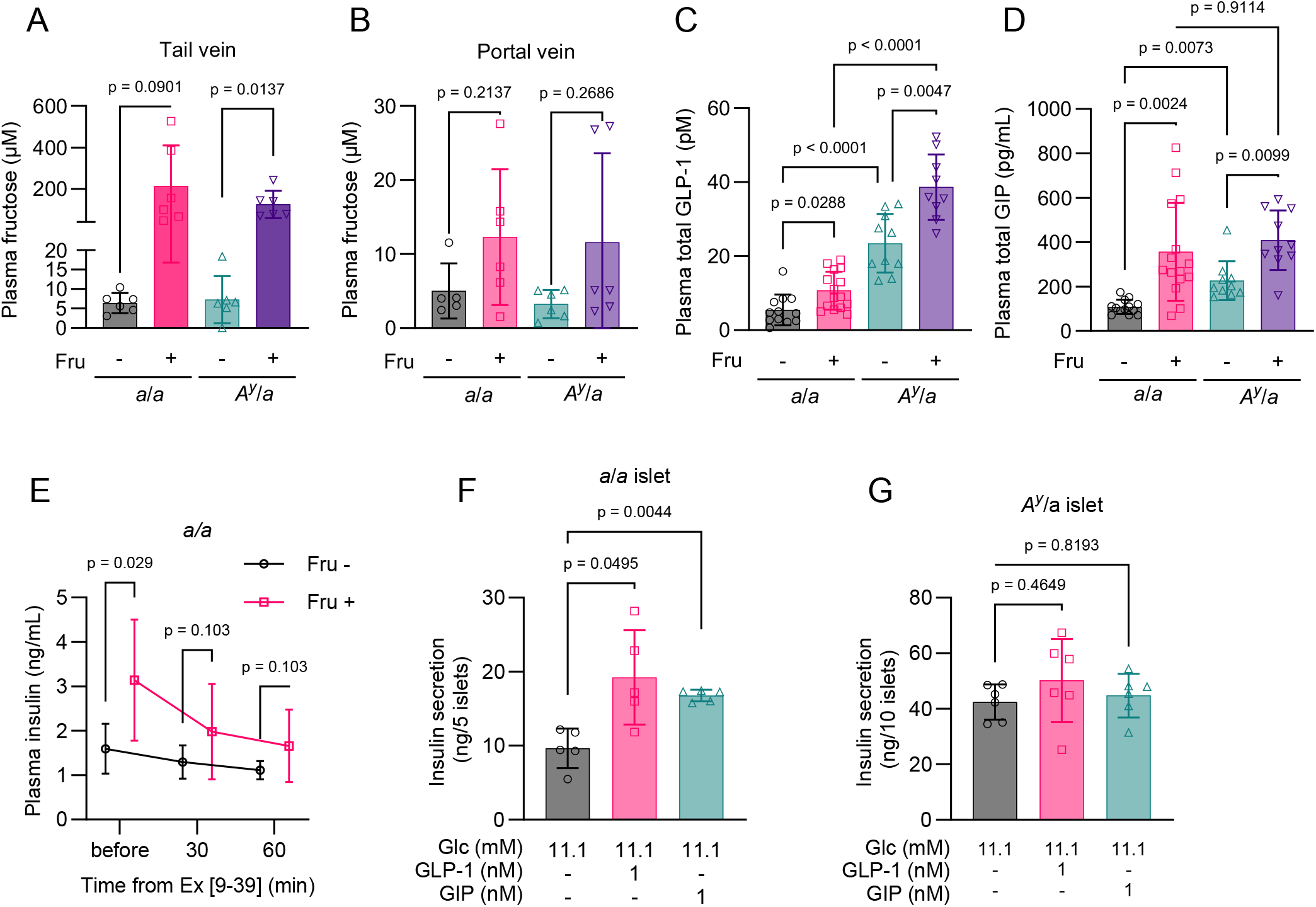
Involvement of plasma fructose, GLP-1, and GIP in fructose-induced plasma insulin response. A. –D. Plasma concentrations of fructose, total GLP-1, and total GIP following 24-hour fructose ingestion. A. Fructose levels in the tail vein. *a*/*a* Fru −: n = 6; *a*/*a* Fru +: n = 6; *A*^*y*^/*a* Fru −: n = 6; *A*^*y*^/*a* Fru +: n = 6. N = 2. B, Fructose levels in the portal vein. *a*/*a* Fru −: n = 5; *a*/*a* Fru +: n = 6; *A*^*y*^/*a* Fru −: n = 6; *A*^*y*^/*a* Fru +: n = 6. N = 2. C, total GLP-1. *a*/*a* Fru −: n = 12; *a*/*a* Fru +: n = 15; *A*^*y*^/*a* Fru −: n = 10; *A*^*y*^/*a* Fru +: n = 9. N = 2. D, total GLP-1. *a*/*a* Fru −: n = 13; *a*/*a* Fru +: n = 15; *A*^*y*^/*a* Fru −: n = 10; *A*^*y*^/*a* Fru +: n = 10. N = 2. E. Effect of exendin [9-39] on fructose-induced insulin response. Exendin [9-39] (100 nmol/kg) was intraperitoneally injected following 24-hour fructose ingestion. Time-dependent changes in plasma insulin levels before and after injection are shown. n = 9, N = 2. F. –G. Insulin secretory responses to GLP-1 and GIP in isolated islets. F, *a*/*a* mouse islets. n = 5. Islets were pooled from two mice. G, *A*^*y*^/*a* mouse islets. n = 6. Islets were pooled from two mice. N = 2. Data are presented as mean ± standard deviation (SD). Fru −: water-administered mice, Fru +: fructose-administered mice. Statistical comparisons were performed using Welch’s one-way ANOVA with Dunnett’s post-hoc test except for E. Welch’s unpaired two-tailed t-test was used for E.

Therefore, our investigation focused on insulinotropic hormones, including GLP-1, GIP, and glucagon. In the fasting state without fructose, *A*^*y*^/*a* mice exhibited elevated GLP-1 and GIP levels compared to *a*/*a* mice (Fig. 2, C–D). Fructose administration significantly increased GLP-1 and GIP levels in both *a*/*a* and *A*^*y*^/*a* mice (Fig. 2, C–D). Given that GLP-1 values are known to vary with different assay systems (Windeløv et al., 2017), we verified our findings using different assay previously validated for similar experiments (Kuhre et al., 2014). The results showed a consistent trend across the four groups, confirming the reliability of the assay system (Fig. A2C and Fig. 2C). Glucagon levels remained unaltered by fructose (Fig. A2D). Interestingly, exendin [9–39], a GLP-1 receptor antagonist, blocked the fructose-induced increase in insulin levels in *a*/*a* mice (Fig. 2E), indicating that GLP-1 is required for the insulinotropic effect of fructose. We then asked why GLP-1 and GIP failed to increase insulin levels in *A*^*y*^/*a* mice. In isolated *a*/*a* islets, both GLP-1 and GIP potentiated insulin secretion (Fig. 2F). In contrast, *A*^*y*^/*a* islets showed high insulin secretion at 11.1 mM glucose alone, with no further increase upon GLP-1 or GIP (Fig. 2G). This pattern is characteristic of mouse β-cells under metabolic overload, which exhibit exaggerated secretion at basal to mid-range glucose but blunted responses to additional secretagogues (Hudish et al., 2019). These findings indicate that compromised β-cell function limits fructose-induced insulin response in diabetes, despite preserved GLP-1 secretion.

To better understand the roles of GLP-1 and GIP in fructose-induced insulin response, we studied glucagon knockout mice (*Gcg*−/−) (Hayashi et al., 2009) and GIP receptor knockout mice (*Gipr*−/−) (Miyawaki et al., 1999) and compared them to their background strain B6. Initially, in B6 mice, fructose administration led to a slight, non-significant increase in blood glucose levels (Fig. 3A), accompanied by increases in insulin, GLP-1, and GIP levels (Fig. 3, B–D). Next, *Gcg*−/−mice, which lack proglucagon-derived peptides, such as GLP-1 and glucagon, due to a *gfp* insertion that disrupts the proglucagon gene (Hayashi et al., 2009), showed significantly higher blood glucose levels after fructose administration (Fig. 3A), whereas insulin and GIP levels remained unchanged (Fig. 3, B and D). These findings suggest that the insulin response to fructose is defective when both GLP-1 and GIP responses are absent, although the reason for the lack of a GIP response in *Gcg*−/−mice remains unclear. Lastly, *Gipr*−/−mice exhibited fructose-induced increases in insulin, GLP-1, and GIP levels (Fig. 3, B–D), with no rise in blood glucose levels (Fig. 3A), phenocopying B6 mice. Thus, the insulin response to fructose remained intact, even without the GIP response. Collectively, these findings suggest that GIP is nonessential, whereas GLP-1 is required for the fructose-induced insulin response. This observation aligns with the exendin [9-39] result (Fig. 2E).

**Figure 3.**
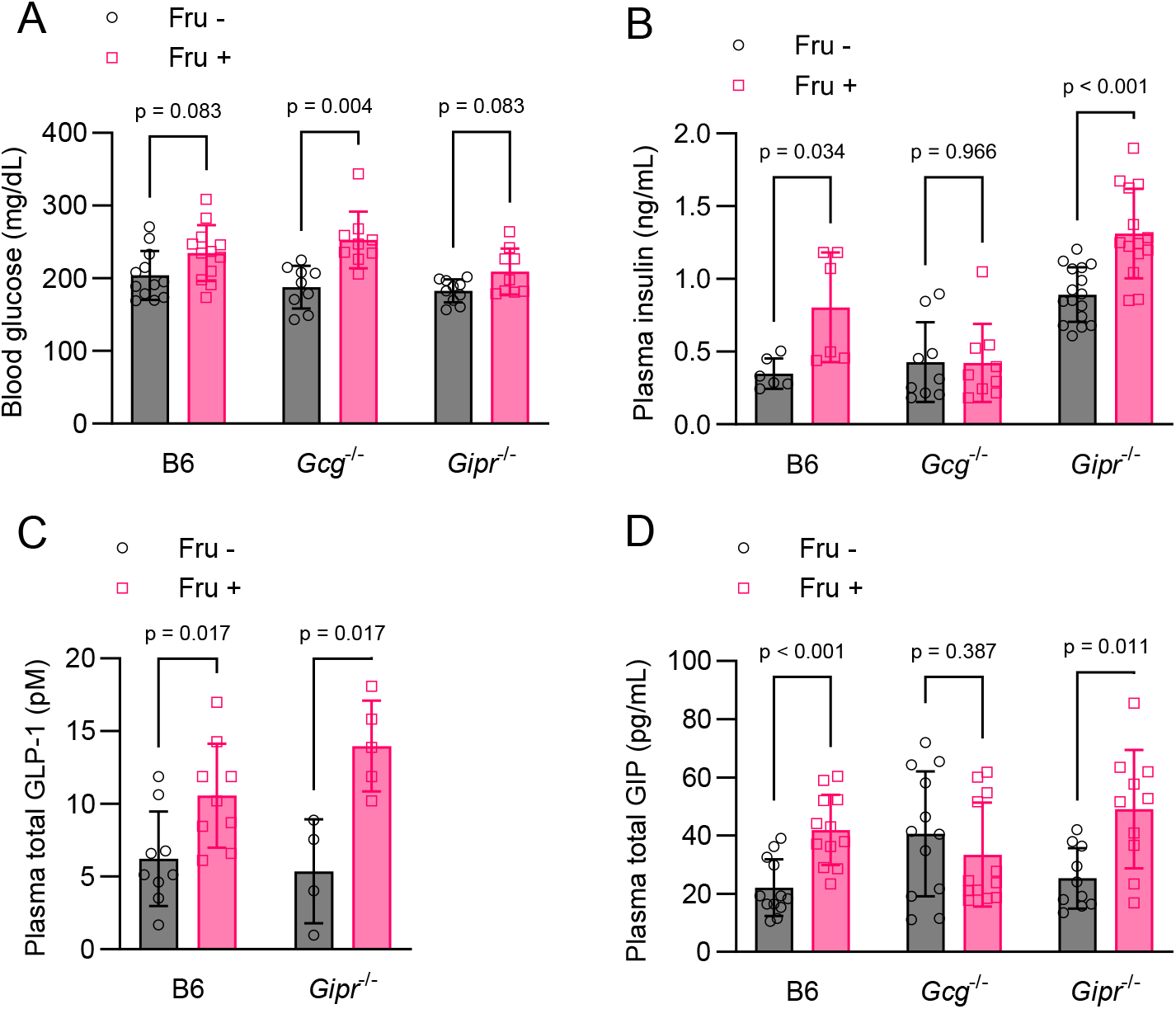
Involvement of GLP-1 and GIP in fructose-induced metabolic responses. Metabolic profiles of B6, *Gcg*^−/−^, and *Gipr*^−/−^ mice following 24-hour fructose ingestion. A. Blood glucose, B6 Fru −: n = 12; B6 Fru +: n = 12; *Gcg*^−/−^ Fru −: n = 9; *Gcg*^−/−^ Fru +: n = 9; *Gipr*^−/−^ Fru −: n = 10; *Gipr*^−/−^ Fru +: n = 9. B. Insulin, B6 Fru −: n = 6; B6 Fru +: n = 6; *Gcg*^−/−^ Fru −: n = 9; *Gcg*^−/−^ Fru +: n = 9; *Gipr*^−/−^ Fru −: n = 16; *Gipr*^−/−^ Fru +: n = 14. C. Total GLP-1, B6 Fru −: n = 9; B6 Fru +: n = 9; *Gipr*^−/−^ Fru −: n = 4; *Gipr*^−/−^ Fru +: n = 5. D. Total GIP, B6 Fru −: n = 12; B6 Fru +: n = 12; *Gcg*^−/−^ Fru −: n = 12; *Gcg*^−/−^ Fru +: n = 12; *Gipr*^−/−^ Fru −: n = 10; *Gipr*^−/−^ Fru +: n = 10. Data are presented as mean ± standard deviation (SD). Statistical comparisons were performed using Welch’s unpaired two-tailed t-test.

### 5.3. GLP-1 secretion is coupled to intracellular fructose metabolism

We explored the mechanisms by which ingested fructose enhanced GLP-1 secretion. Earlier studies have suggested mechanisms similar to GIIS: fructose metabolism produces ATP in L-cells, causing K_ATP_ channels to close, which leads to membrane depolarisation and subsequent Ca^2+^ entry (Kuhre et al., 2014; Gribble et al., 2003). This theory is supported by several GLUTag cell experiments; however, further validation is warranted, as noted in the Discussion section. Our aim was to confirm whether fructose metabolism increases the ATP/ADP ratio, promotes Ca^2+^ influx, and stimulates GLP-1 secretion.

Fructose dose-dependently enhanced GLP-1 secretion at concentrations above 10 mM (Fig. 4A). To investigate the involvement of fructose metabolism, we used siRNA to knock down two key enzymes involved in fructolysis: ketohexokinase (*Khk*) and aldolase B (*Aldob*) (Fig. A3, A–B). Intracellular ATP/ADP ratio and calcium ([Ca^2+^]i) were visualized using a Perceval HR biosensor (Tantama et al., 2013) and Fluo-4, respectively. Fructose triggered a gradual increase in the ATP/ADP ratio in control cells treated with non-targeting siRNA (siNT). This elevation was inhibited when *Khk* and *Aldob* were depleted (Fig. 4, B–C, Fig. A3, C–D). Similarly, fructose triggered a gradual increase in [Ca^2+^] i with successive firing in siNT cells (Fig. 4D). Knockdown of *Khk* and *Aldob* inhibited [Ca^2+^]i firing (Fig. 4D) and reduced average [Ca^2+^]i responses (Fig. A3E), as quantified by the incremental area under the curve (iAUC) (Fig. A3F). Interestingly, both the ATP/ADP ratio and [Ca^2+^]i exhibited similar time-dependent trends, initially experiencing a transient decline upon fructose stimulation, followed by a steady rise. This initial decline might be attributed to swift ATP utilisation by KHK, as previously observed in enterocytes (Merino et al., 2019), whereas the subsequent increase could be linked to ATP replenishment through fructose catabolism.

**Figure 4.**
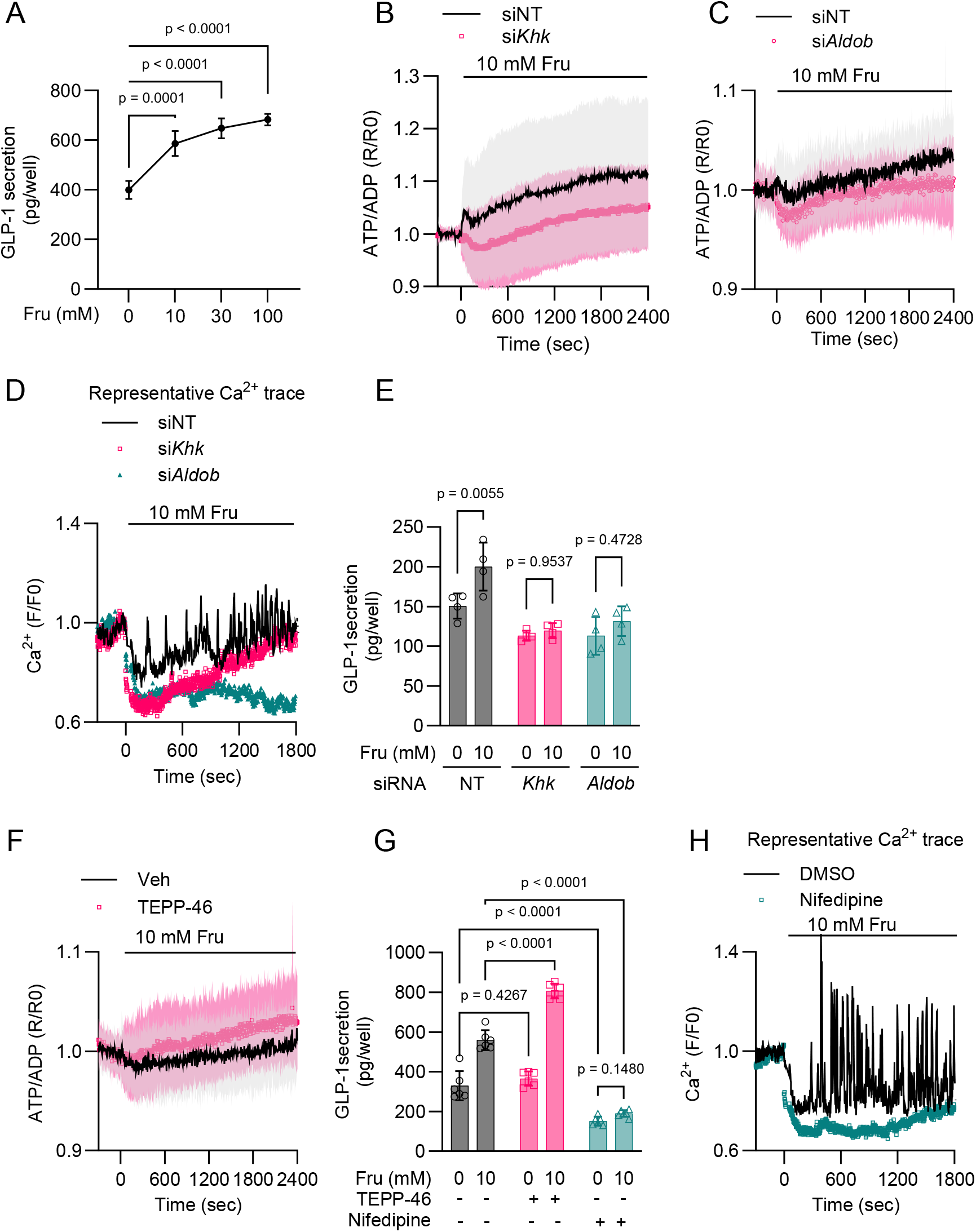
Metabolism-secretion coupling in fructose-induced GLP-1 secretion in GLUTag cells. A. Dose-dependent effect of fructose on GLP-1 secretion. n = 6. N = 3. B. –C. Effect of fructose on the intracellular ATP/ADP ratio, as visualised by Perceval-HR. Effects of *Khk* and *Aldob* knockdown were tested, and the time course of the normalised ratiometric signals (R/R0) is indicated. For statistical analyses, see Fig. A3, C–D. B. siNT (solid lines): n = 26; si*Khk* (circles): n = 38. N = 2. C. siNT (solid lines): n = 26; si*Aldob* (circles): n = 27. N = 2. D. Effect of fructose on intracellular Ca^2+^ was visualised by Fluo-4. Symbols: siNT, solid lines; si*Khk*, squares; si*Aldob*, upward triangles. Representative traces of normalised fluorescence (F/F0) are shown. For statistical analyses, see Fig. A3, E–F. N = 2. E. Effects of *Khk* and *Aldob* knockdown on fructose-induced GLP-1 secretion. n = 4. N = 2. F. Effect of TEPP-46 (10 μM) on fructose-induced increase in the intracellular ATP/ADP ratio, as visualised by Perceval-HR. Veh (solid lines): n = 32; TEPP-46 (squares): n = 45. The time course of the normalised ratiometric signals (R/R0) is shown. For statistical analyses, see Fig. A3G. N = 2. G. Effects of TEPP-46 (10 μM) and nifedipine (10 μM) on fructose-induced GLP-1 secretion. n = 6. N = 2. H. Effect of nifedipine (10 μM) on fructose-induced Ca^2+^ response as visualised by Fluo-4. Representative traces of normalised fluorescence (F/F0) are shown. For statistical analysis, see Fig. A3, H–I. N = 2. Symbols: DMSO, solid lines; nifedipine, squares. All experiments were performed using GLUTag cells. For imaging experiments, 10 mM fructose was loaded at t = 0. siNT, non-targeting siRNA. Data are presented as mean ± SD. Statistical comparisons were performed using Welch’s one-way ANOVA with Dunnett’s post-hoc test for A, and two-way ANOVA with Šídák’s multiple comparisons post-hoc test for E and G.

These knockdown cells exhibited impaired fructose-induced GLP-1 secretion (Fig. 4E). These findings indicate that the suppression of fructose metabolism leads to a decrease in the ATP/ADP ratio, [Ca^2+^]i response, and GLP-1 secretion. Research has shown that TEPP-46, an activator of pyruvate kinase M2 (PKM2), enhances fructose metabolism in both intestinal epithelial cells (Taylor et al., 2021) and β-cells (Murao et al., 2025). These findings prompted us to use TEPP-46 to confirm the link between metabolism and secretion. In GLUTag cells, TEPP-46 significantly enhanced fructose-induced elevation of the ATP/ADP ratio (Fig. 4F, Fig. A3G), and GLP-1 secretion (Fig. 4G). Fructose-induced [Ca^2+^]i response and GLP-1 secretion were suppressed by nifedipine, an inhibitor of L-type voltage-dependent Ca^2+^ channels (VDCCs) (Fig. 4, G–H), indicating that Ca^2+^ influx through L-type VDCC plays a permissive role in fructose-induced GLP-1 secretion. Collectively, these findings demonstrate that fructose metabolism increases the ATP/ADP ratio, which is positively correlated with [Ca^2+^]i and GLP-1 secretion.

### 5.4. Functional K_ATP_ channels are required for Fructose-induced GLP-1 secretion in vivo

To validate the involvement of K_ATP_ channels in fructose-induced GLP-1 secretion in vivo, we utilised *Kcnj11*^−/−^ mice, which lack K_ATP_ channel activity throughout the body due to disruption of the pore-forming subunit KCNJ11 (Miki et al., 1998). Wild-type littermates (*Kcnj11*^+/+^) displayed a phenotype comparable to that of B6 mice (Fig. 5, A–D). In contrast, *Kcnj11*^−/−^ mice exhibited null fructose-induced insulin and GLP-1 responses (Fig. 5, B–C), resulting in a significant increase in glycaemia (Fig. 5A).

**Figure 5.**
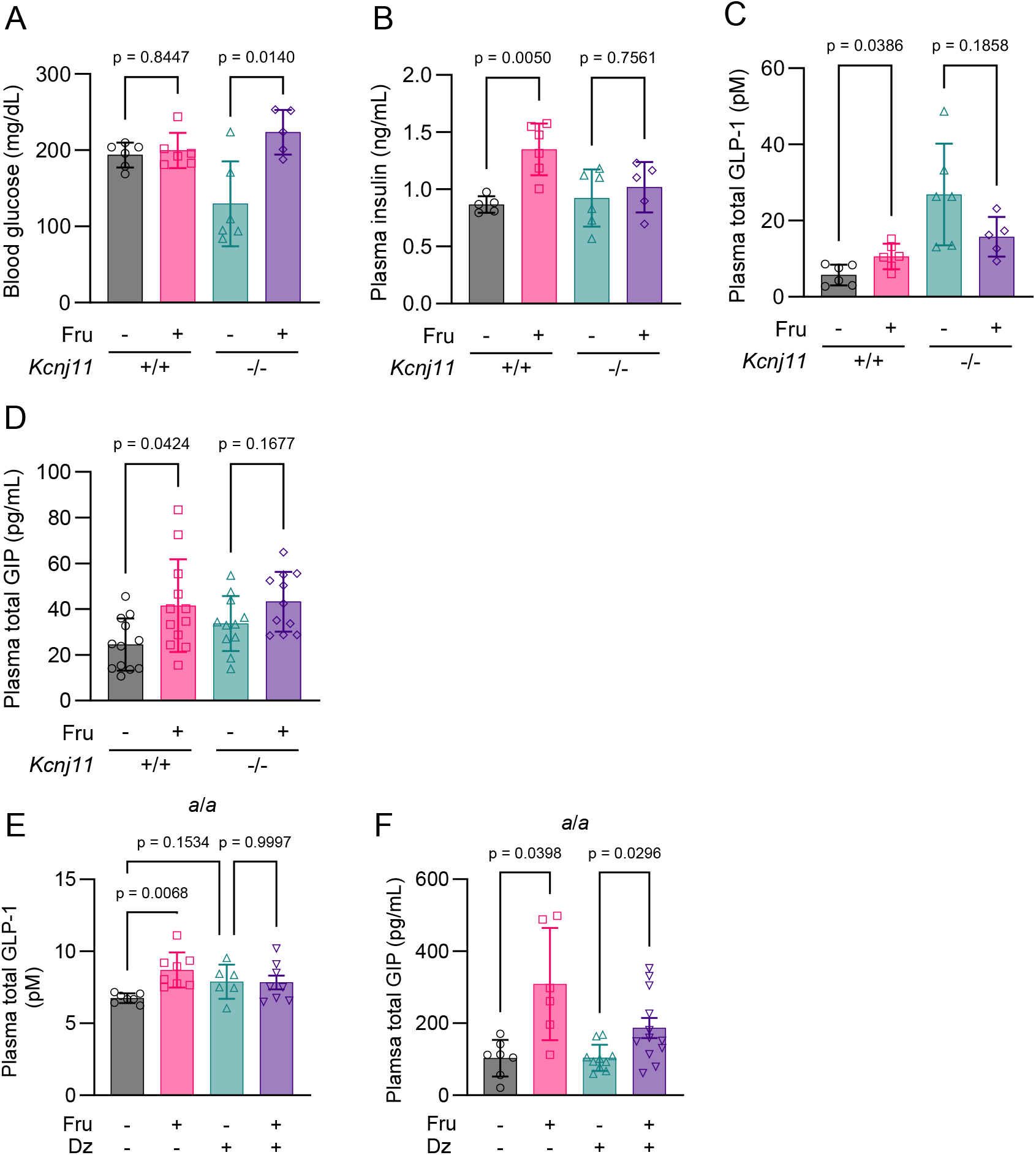
Involvement of K_ATP_ channels in fructose-induced GLP-1 and GIP secretion in vivo. A. –D. Metabolic profiles of *Kcnj11*^*+*/+^ and *Kcnj11*^−/−^ mice following 24-hour fructose ingestion. N = 2. A. Blood glucose, −/− Fru −: n = 6; −/− Fru −: n = 6; −/− Fru −: n = 6; −/− Fru −: n = 5. B. Insulin, −/− Fru −: n = 5; −/− Fru −: n = 6; −/− Fru −: n = 6; −/− Fru −: n = 5. C. Total GLP-1, −/− Fru −: n = 5; −/− Fru −: n = 6; −/− Fru −: n = 6; −/− Fru −: n = 5. D. Total GIP, −/− Fru −: n = 12; −/− Fru −: n = 12; −/− Fru −: n = 12; −/− Fru −: n = 11. E. –F. Effect of diazoxide (Dz) on fructose-induced GLP-1 and GIP responses in *a*/*a* mice. Diazoxide (10 mM) was added to the water or fructose solution. Incretin levels were measured following 24-hour fructose ingestion. N = 2. E. Total GLP-1, Fru − Dz −: n = 7; Fru + Dz −: n = 8; Fru − Dz +: n = 6; Fru + Dz +: n = 7. F. Total GIP, Fru − Dz −: n = 7; Fru + Dz −: n = 6; Fru − Dz +: n = 10; Fru + Dz +: n = 12. Data are presented as mean ± SD. Statistical comparisons were performed using Welch’s unpaired two-tailed t-test for A–D, and Welch’s one-way ANOVA with Dunnett’s post-hoc test for E–F.

Furthermore, administration of diazoxide, a K_ATP_ channel opener, inhibited the fructose-induced increase in GLP-1, but not GIP, in *a*/*a* mice (Fig. 5, E–F). These findings demonstrate that fructose-induced GLP-1 secretion is dependent on functional K_ATP_ channels in vivo.

### 5.5 Characterization of Fructose metabolism in lean and diabetic mouse intestinal crypts

GLP-1 levels were higher in *A*^*y*^/*a* mice than in *a*/*a* mice at baseline and were further increased by fructose in both strains (Fig. 2C). We investigated whether these changes were due to fructose metabolism in L-cells. Initially, immunohistochemical analysis was performed on L-cells in the jejunum and ileum, which are the primary sites of fructose absorption (Bode et al., 1981). Although most L-cells were located in crypts, KHK was mainly expressed in villi (Fig. 6A). Notably, KHK was also faintly detected in crypt cells, including L-cells, in *A*^*y*^/*a* mice, whereas it was entirely absent in crypts of *a/a* mice (Fig. 6A, see insets), indicating increased KHK expression in *A*^*y*^/*a* L-cells. The density of crypts was similar between the genotypes (Fig. 6B), whereas the density of L-cells was slightly lower in *A*^*y*^/*a* mice (Fig. 6B), suggesting that elevated GLP-1 levels cannot be attributed to an increased number of L-cells.

**Figure 6.**
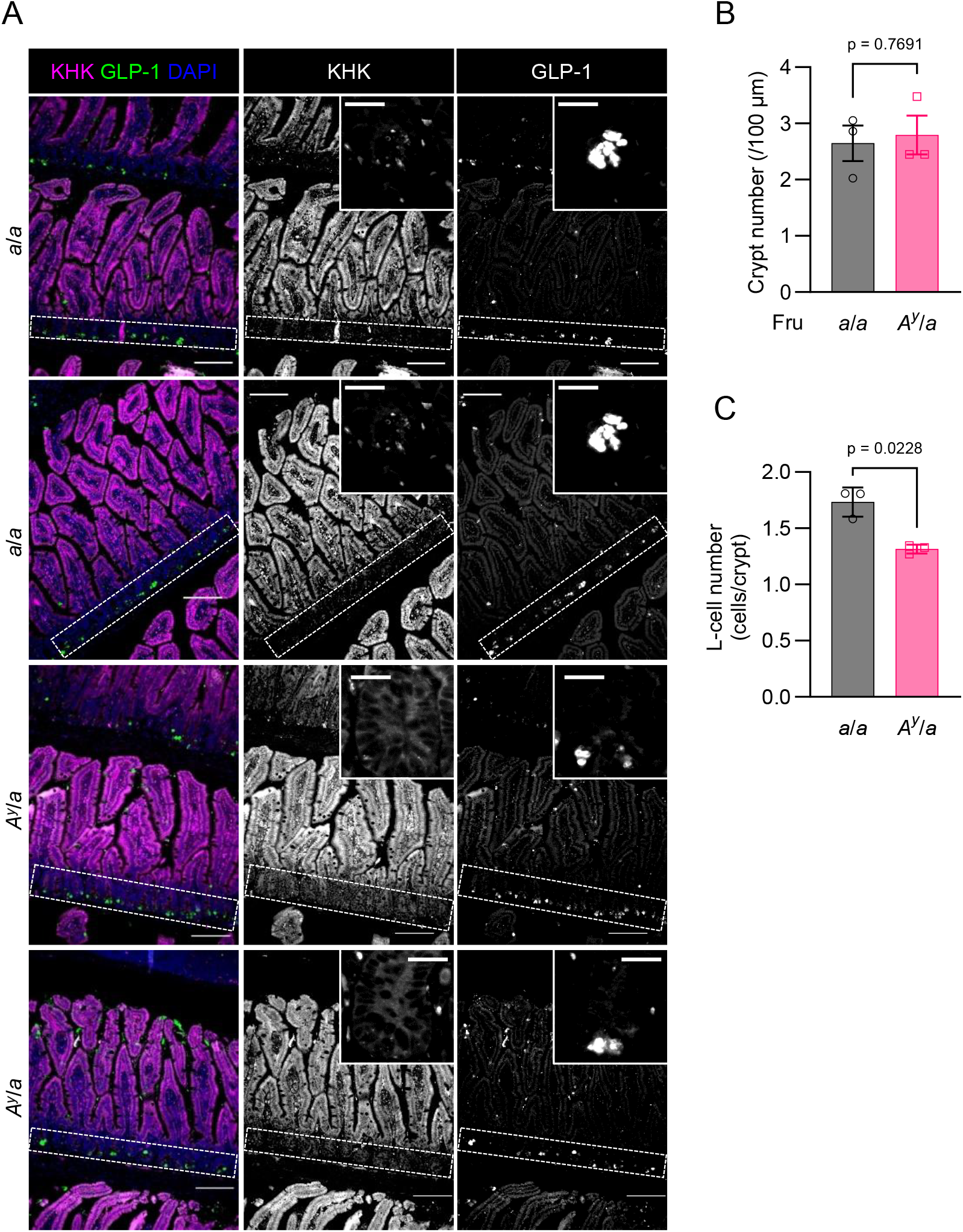
Histology of L-cells in lean and diabetic mice. A. Immunofluorescent images of the ileum from *a*/*a* and *A*^*y*^/*a* mice. Left: Overlay images of DAPI, KHK, and GLP-1. Centre and right: corresponding grayscale images of separate channels for KHK and GLP-1. The dashed rectangles indicate a layer of crypts. The insets in the grayscale panels show representative crypts. Scale bars, main 100 μm, inset 20 μm. n = 2 mice/genotype. N = 2. B. Quantification of crypt number normalised to mucosal length. n = 3 mice/genotype. Five fields per mouse were pooled into a single value. C. Quantification of L-cell numbers normalised to crypt numbers. n = 3 mice/genotype. Five fields per mouse were pooled into a single value. Data are presented as mean ± standard deviation (SD). Statistical comparisons were performed using Welch’s unpaired two-tailed t-test.

Given this, we aimed to quantify the metabolic activity of fructose in these cells. Purified L-cells were technically unsuitable for metabolomic analysis owing to the scarcity of these cells after sorting. As an alternative strategy, we used isolated intestinal crypts. Initially, the isolation procedure was verified using *Gcg*^gfp/+^ mouse crypts, in which GFP-positive L-cells were successfully identified (Fig. A4A). Crypts from *a*/*a* and *A*^*y*^/*a* mice were exposed to 50 mM ^13^C_6_-fructose, a stable isotopomer in which all six carbon atoms are ^13^C. The potential metabolic pathways of ^13^C_6_-fructose are shown in Fig. 7A. Isotopomers with 0-6 carbon atoms substituted by ^13^C are denoted as M-M6. Naturally occurring isotopomers are represented by M and M1. The ^13^C_6_-fructose can be metabolised into F1P [M6] (fructose 1-phosphate), GA3P [M3] (glyceraldehyde 3-phosphate), and DHAP [M3] (dihydroxyacetone phosphate). Subsequently, GA3P [M3] enters glycolysis and the TCA cycle, resulting in the formation of malate [M2–M4] and citrate [M2–M6]. These metabolites contribute to ATP generation through oxidative phosphorylation (OXPHOS). Another metabolic route for GA3P [M3] is its conversion to lactate [M3], which produces ATP. Additionally, GA3P [M3] and DHAP [M3] can be utilised in gluconeogenic processes to synthesise M6 G6P.

**Figure 7.**
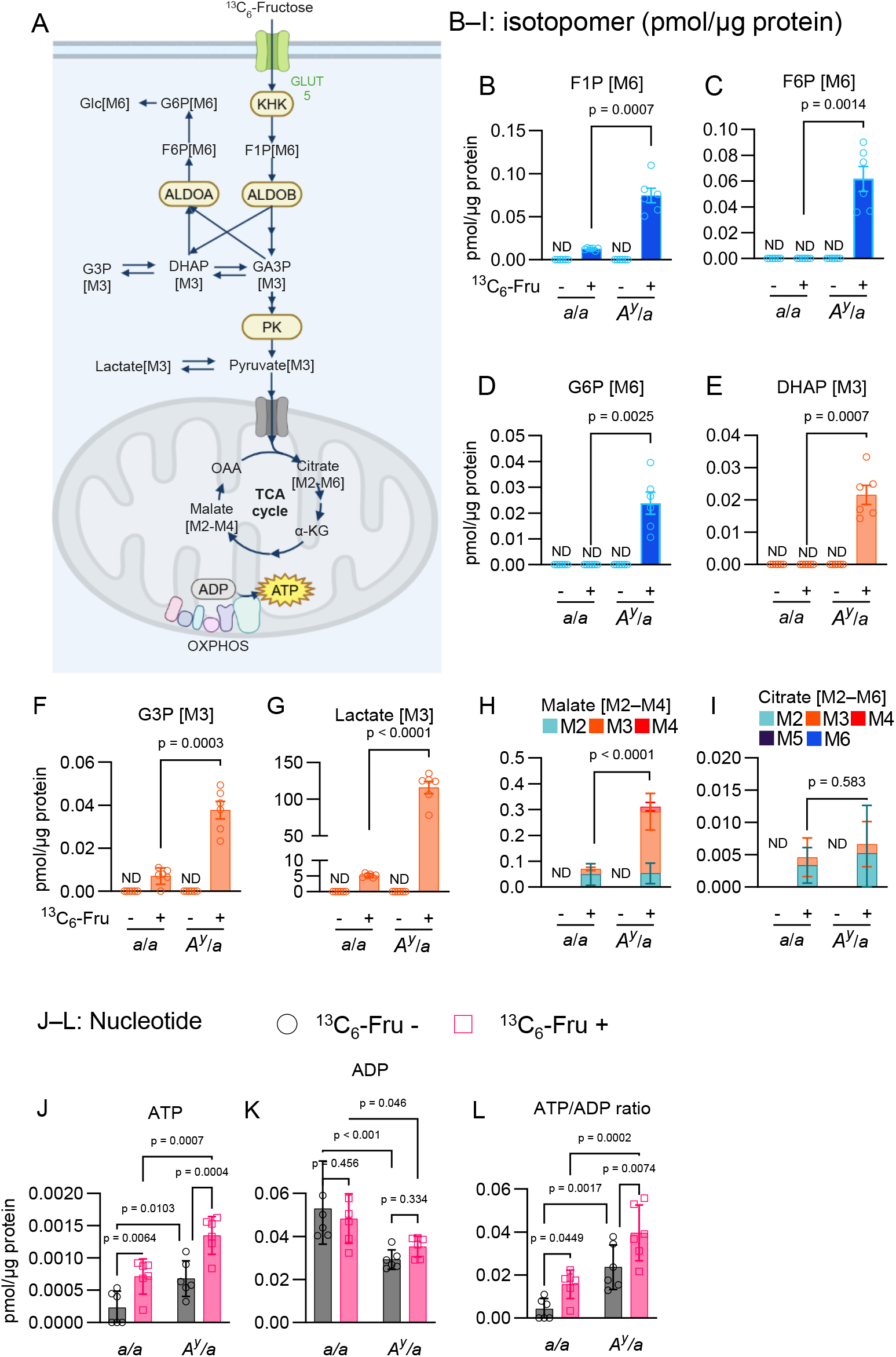
Fructose metabolic tracing in isolated intestinal crypts of lean and diabetic mice. A. Schematic overview of the potential metabolic fate of ^13^C_6_-fructose in the intestine. G6P, glucose 6-phosphate; F6P, fructose 6-phosphate; F1P, fructose 1-phosphate; GA3P, glyceraldehyde 3-phosphate; DHAP, dihydroxyacetone phosphate; G3P, glycerol 3-phosphate; OXPHOS, oxidative phosphorylation. M2–M6 denote isotopomers with 0-6 carbon atoms substituted by ^13^C. Drawn using BioRender.com. B. –I. ^13^C_6_-fructose tracing in intestinal crypts isolated from *a*/*a* and *A*^*y*^/*a* mice. Only Isotopomers converted from ^13^C_6_-fructose are indicated. See Fig. A4B for comprehensive results, including those of other isotopomers. n = 6 for each condition. N = 2. J. –L. Adenosine nucleotide content in ^13^C_6_-fructose tracing. Only unlabelled isotopomers [M0] were analysed. n = 6 for each condition. N = 2. Data are presented as mean ± SD. ND, not detected. Statistical comparisons were performed using Welch’s unpaired two-tailed t-test for B–I and two-way ANOVA with Uncorrected Fisher’s Least Significant Difference test for J–L. For H and I, the sum of all the indicated isotopomers was compared.

Fig. 7, B–I show the quantity of ^13^C_6_-Fru-derived isotopomers in *a*/*a* and *A*^*y*^/*a* mouse crypts in the presence or absence of ^13^C_6_-Fru. As anticipated, these isotopomers were detected in stimulated (^13^C_6_-Fru +) crypts but not in unstimulated (^13^C_6_-Fru −) crypts. F1P [M6], the most upstream intermediate, was significantly upregulated in ^13^C_6_-Fru + *A*^*y*^/*a* crypts compared to ^13^C_6_-Fru + *a*/*a* crypts (Fig. 7B). All downstream isotopomers, except citrate [M2–M6], were also upregulated in ^13^C_6_-Fru + *A*^*y*^/*a* crypts. Collectively, these results suggest that increased KHK expression leads to an enhanced flux into fructolysis in *A*^*y*^/*a* mouse crypts. Fig. 7, J–L indicate the quantity of unlabelled ATP and ADP and their ratio in the crypts. In the absence of ^13^C_6_-Fru, ATP levels were elevated and ADP levels were decreased, resulting in an increased ATP/ADP ratio in *A*^*y*^/*a* crypts compared to *a*/*a* crypts. ATP levels and the ATP/ADP ratio were further increased by ^13^C_6_-Fru stimulation in both genotypes. These trends in ATP and ATP/ADP ratio were consistent with the differential GLP-1 levels in basal and fructose-stimulated states (Fig. 2C), suggesting that the metabolic status of the intestine is reflected in fructose-induced GLP-1 secretion in vivo.

## 6. DISCUSSION

This study delineates the mechanism linking short-term fructose consumption to glycaemic control: (1) fructose stimulates GLP-1 secretion via intestinal fructose metabolism that elevates the ATP/ADP ratio and closes K_ATP_ channels, and (2) GLP-1 then potentiates insulin secretion, thereby suppressing fructose-induced hyperglycaemia.

Fructose modulates glycaemia through multiple mechanisms operating on different timescales. First, acute fructose administration can transiently increase blood glucose levels (Kuhre et al., 2014; Seino et al., 2015; Kong et al., 1999). This is often interpreted to suggest that the insulin response is simply a secondary effect of glycaemia. However, we observed a robust insulin increase in *a*/*a* mice, without a concomitant change in blood glucose (Fig. 1, B–C), arguing against a secondary mechanism. The early glycaemic increase is at least partly attributable to intestinal fructose-to-glucose conversion (Ockerman and Lundborg, 1965; Jang et al., 2018). Consistent with this, triose/hexose-phosphate intermediates (DHAP [M3], F6P [M6], and G6P [M6]) were substantially increased in *A*^*y*^/*a* crypts (Fig. 7, C–E), suggesting enhanced intestinal conversion as a contributor to the larger glycaemic excursion in *A*^*y*^/*a* mice (Fig. 1B).

Second, prolonged fructose exposure induces hepatic insulin resistance (Softic et al., 2020; Hannou et al., 2018), typically after weeks of intake (Basciano et al., 2005; Ter Horst et al., 2016). Mechanistically, fructose stimulates hepatic glucose production by upregulating glucose-6-phosphatase (G6PC) via carbohydrate response element–binding protein (ChREBP), partly overriding insulin action (Kim et al., 2019). Whether this has contributed to fructose-induced hyperglycaemia in our setting is uncertain. However, two observations argued against a dominant role: fructose ingestion lowered OGTT excursions in lean mice (Fig. 1D), suggesting that insulin-mediated suppression of endogenous glucose production was preserved. Fructose-induced hyperglycaemia was greatest in strains lacking an insulin response (ΔBG: *A*^*y*^/*a* 80.17; *Gcg*^−/−^ 64.78; *Kcnj11*^−/−^ 93.93) compared with strains with intact responses (ΔBG: *a*/*a* 12.30; B6 30.58; *Gipr*^−/−^ 26.62). Together, these findings indicate that fructose-evoked insulin increments are effective in restraining glycaemia, whereas failure of this response permits larger glycaemic excursions.

As a potential mechanism for fructose-induced insulin response, in vitro studies have demonstrated that high concentrations (>10 mM) of fructose can directly potentiate GIIS (Grodsky et al., 1963; Ashcroft et al., 1972; Zawalich et al., 1977; Grant et al., 1980; Murao et al., 2025). However, these effects have not been substantiated in vivo, likely because plasma fructose levels fall far below the concentration required for the direct stimulation of β-cells. Indeed, the observed plasma fructose levels (~100 μM) were in good agreement with previous measurements using 20% fructose diet-fed B6 mice (Patel et al., 2015) and sucrose-administered Wistar rats (Sugimoto et al., 2010). Human plasma fructose levels are reported to be below 100 μM in the postprandial state (Kawasaki et al., 2004).

Glucose induces depolarising currents in L-cells via two mechanisms: (1) SGLT-mediated Na^+^/glucose transport and (2) GLUT2-mediated glucose uptake and intracellular metabolism, leading to K_ATP_ channel closure (Reimann et al., 2008; Kuhre et al., 2015; Hjørne et al., 2022). Both mechanisms lead to Ca^2+^ influx through VDCCs, which was demonstrated through its inhibition by nifedipine in GLUTag cells (Reimann et al., 2005), perfused rat intestines (Kuhre et al., 2015), and human intestine fragments (Sun et al., 2017).

As fructose is not a substrate for SGLT1 (Koepsell, 2020), it has been speculated that fructose-induced GLP-1 secretion relies solely on intracellular fructose metabolism and K_ATP_ channel closure. This hypothesis has been corroborated by several findings derived from GLUTag cells, including fructose-induced K_ATP_ channel closure and action potentials (Gribble et al., 2003), fructose-induced increases in intracellular NADPH levels (Kuhre et al., 2014), and inhibition of fructose-induced GLP-1 secretion by diazoxide (Kuhre et al., 2014). However, prior studies lacked evidence regarding whether fructose metabolism elevates the ATP/ADP ratio and [Ca^2+^]i, which we demonstrated in GLUTag cells (Fig. 4). Additionally, the link between fructolysis activity, ATP/ADP ratio, and K_ATP_ channels was validated both in vivo and ex vivo in *Kcnj11*^−/−^ mice (Fig. 5) and intestinal crypts (Fig. 7), suggesting that these results are not limited to a specific cell line. In contrast to the current study, our previous research showed that a substantial GLP-1 response occurs even in *Kcnj11*^−/−^ mice 15 min after gavage with excess fructose (6 g/kg) (Seino et al., 2015). Potentially, fructose-induced GLP-1 response involves distinct mechanisms in the acute and subacute phases, with the former being at least partially independent of fructose metabolism. Given the substantial volume of liquid administered, mechanical stimuli may contribute to the acute GLP-1 response (Huang et al., 2024).

The effect of fructose ingestion on GIP secretion has been less extensively investigated than that of GLP-1. Fructose can enhance GIP secretion from primary-cultured K-cells (Parker et al., 2009). Consistently, the present study demonstrated that 24-h fructose ingestion robustly increased GIP levels across multiple strains. In contrast, research has shown that acute fructose administration failed to elevate GIP levels in normal mice, rats, and healthy humans (Kuhre et al., 2014). We previously observed that acute fructose ingestion can elicit a GIP response in streptozotocin-induced diabetic mice, but not in normal mice (Seino et al., 2015). These highly variable outcomes suggest that fructose-induced GIP secretion is sensitive to ingestion conditions and metabolic status. Notably, the impaired GIP response in *Kcnj11*^−/−^ mice (Fig. 5D) indicates that K_ATP_ channels are, at least in part, involved in fructose-induced GIP secretion, consistent with the abundant expression of K_ATP_ channel subunits in primary K-cells (Parker et al., 2009). Unexpectedly, fructose-induced GIP response was also deficient in *Gcg*^−/−^ mice, with elevated basal GIP levels in the absence of fructose (compare Fig. 3, G and K). Although the precise mechanism remains unknown, GLP-1 deficiency may alter K-cell function, leading to compensatory hypersecretion of GIP, as previously suggested (Iida et al., 2016). The mechanism underlying fructose-induced GIP secretion requires further investigation.

Compared to previous studies, which typically used acute excessive doses of fructose (2–6 g/kg body weight), our 24-hour ad-lib fructose ingestion is more physiologically relevant, recapitulating the consumption of sugar-sweetened beverages (SSBs). SSBs are a major source of fructose in contemporary diets containing 5–10% fructose (Ando et al., 2023) and are particularly detrimental to glycaemic control (Stanhope et al., 2009; Imamura et al. 2015; Choo et al. 2018; Drouin-Chartier et al. 2019).

Taken together, we propose the following roles for the GLP-1/β-cell axis in fructose-associated metabolic dysfunction. In the healthy state, the GLP-1/β-cell axis counteracts the acute glycaemic effect of fructose, as observed in lean mice. With prolonged fructose exposure, for example habitual sugar-sweetened beverage intake, either independently or through promotion of obesity, the GLP-1/β-cell axis becomes upregulated, leading to chronic hyperinsulinaemia. This view is supported by clinical data showing more pronounced GLP-1 and insulin responses to fructose in obese subjects than in lean subjects (Galderisi et al., 2019). Upregulation of KHK and enhanced fructolysis in L-cells during obesity may contribute to this amplification, which is consistent with our data (Figs. 6 and 7) and Taylor et al. (2021). Under diabetic conditions, the detrimental glycaemic effect of fructose becomes apparent. The GLP-1/β-cell axis no longer mitigates the acute rise in glycaemia because β cells are less responsive (Fig. 2G), and hepatic insulin resistance further limits control of endogenous glucose production. Although our study did not address long-term outcomes, future work should examine how intestinal fructose metabolism and the GLP-1/β-cell axis evolve during chronic fructose consumption.

In this context, the potential contribution of GLP-1 to fructose-induced liver injury remains unclear. Pharmacological GLP-1 receptor agonists decrease hepatic steatosis in patients with metabolic dysfunction–associated steatohepatitis (MASH) (Sanyal et al., 2025) and fructose-fed rodents (Gao et al., 2020). Whether similar benefits arise from the modest nutrient-evoked increase in endogenous GLP-1 is unclear. Mice deficient in the GLP-1 receptor or proglucagon-derived peptides show resistance to hepatic steatosis during high-fat feeding (Ayala et al., 2010; Nishida et al., 2024), which complicates the inference from knockout models. Moreover, DPP-4 inhibitors, which increase active GLP-1 by only a few-fold, have not consistently improved non-alcoholic fatty liver disease (Cui et al., 2016). By analogy, the small GLP-1 increments accompanying fructose ingestion are likely insufficient to counter dominant lipogenic drivers in the liver.

A limitation of this study was the use of intestinal crypts instead of purified L-cells for metabolic analysis. Although various methodologies are available to study L-cells (Kuhre et al., 2021), reliable metabolomic data for L-cells are still lacking. Such an analysis may have been impeded by the difficulty in obtaining a sufficient number of sorted L-cells and by the disruption of the metabolic activity of L-cells through a cell sorting process. One advantage of our approach of maintaining the L-cell resident niche is to preserve L-cell metabolism close to its in situ status. However, the extent to which these results reflect the specific metabolic activity in L-cells remains to be validated. Future technical advancements are warranted for the precise characterisation of L-cell metabolism.

In conclusion, this study identified the intestinal fructose metabolism/GLP-1/β-cell axis as a counter-regulatory mechanism for glycaemia following short-term fructose ingestion. These findings provide significant insights into the systemic fructose metabolism.

## 2. ABBREVIATIONS

iAUC: incremental area under the curve
GIIS: glucose-induced insulin secretion
GIP: glucose-dependent insulinotropic polypeptide
GLP-1: glucagon-like peptide 1
K_ATP_ channel: ATP-sensitive potassium channel
MASH: metabolic dysfunction–associated steatohepatitis
SSB: sugar-sweetened beverage
VDCC: voltage-dependent Ca^2+^ channel

## 7. AUTHOR CONTRIBUTIONS

Conceptualization, N.M.; Methodology, N.M..; Investigation, N.M., R.M., S.H., T.H., E.T., and M.H.; Resources: T.O., N.Y., N.H., and Y.H.; Writing – Original Draft: N.M.; Writing – Review & Editing: N.M., Y.S., Y.Y., Y.H., and A.S.; Data Curation: N.M.; Visualization: N.M.; Supervision: Y.S., Y.Y., and A.S.; Funding Acquisition: N.M.

## 8. ACKNOWLEDGEMENTS

The authors are grateful to President Yutaka Seino (Kansai Electric Power Hospital) for his generous support in this research. The authors thank Professor Daniel Drucker and Dr. Akira Hirasawa for GLUTag cells; Yoshikazu Hoshino (Hoshino Laboratory Animals, Inc) for NSY.B6 mice; Yasuhiro Maeda (Fujita Health University) for assistance with LC-MS. The authors thank Asami Yamaguchi and Megumi Akiyama for their excellent technical assistance.

## 9. DATA AVAILABILITY

Data supporting the findings of this study are available from the corresponding author upon reasonable request.

## 14. APPENDIX: Figs. A1–A4

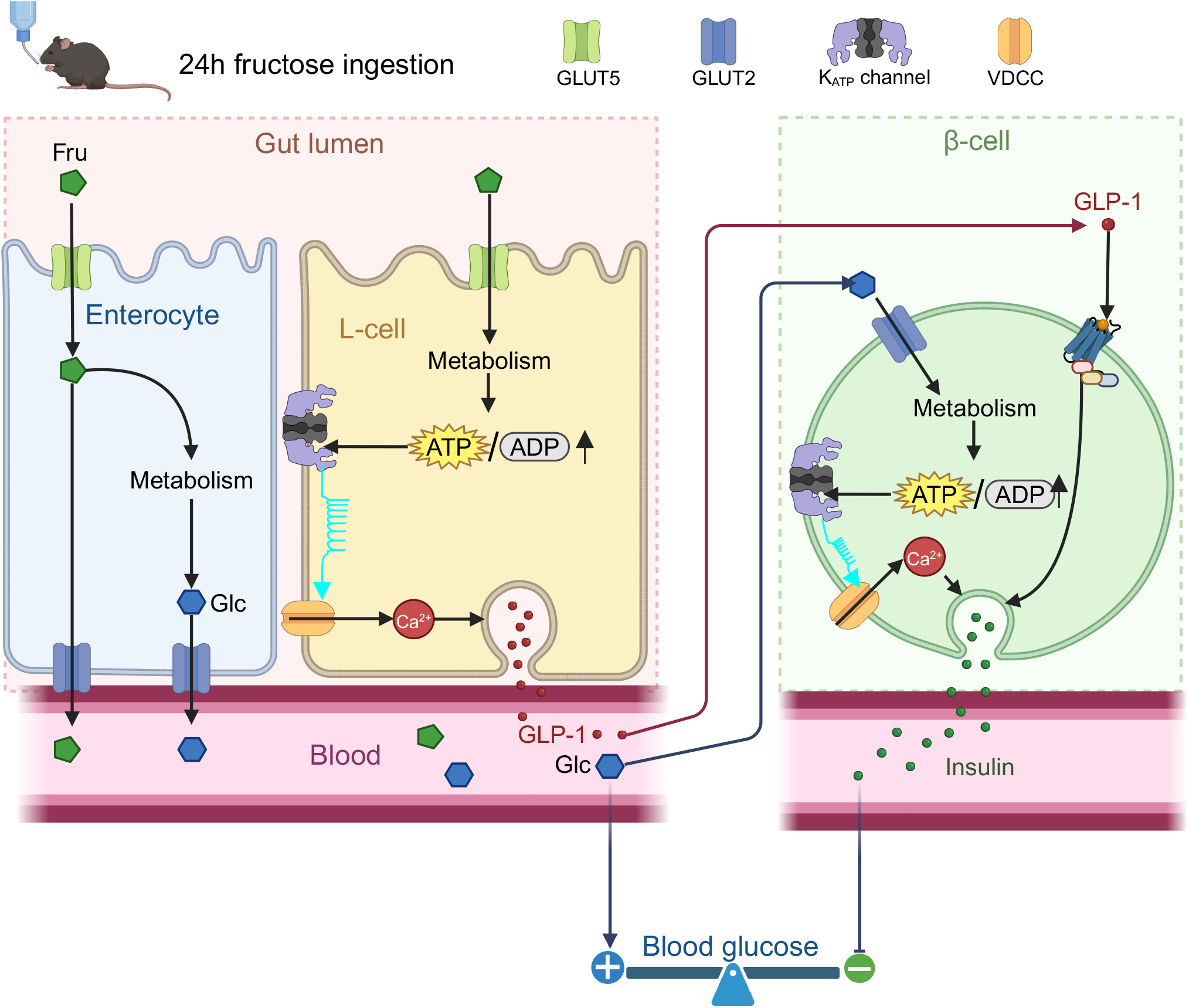

**Figure A1.**
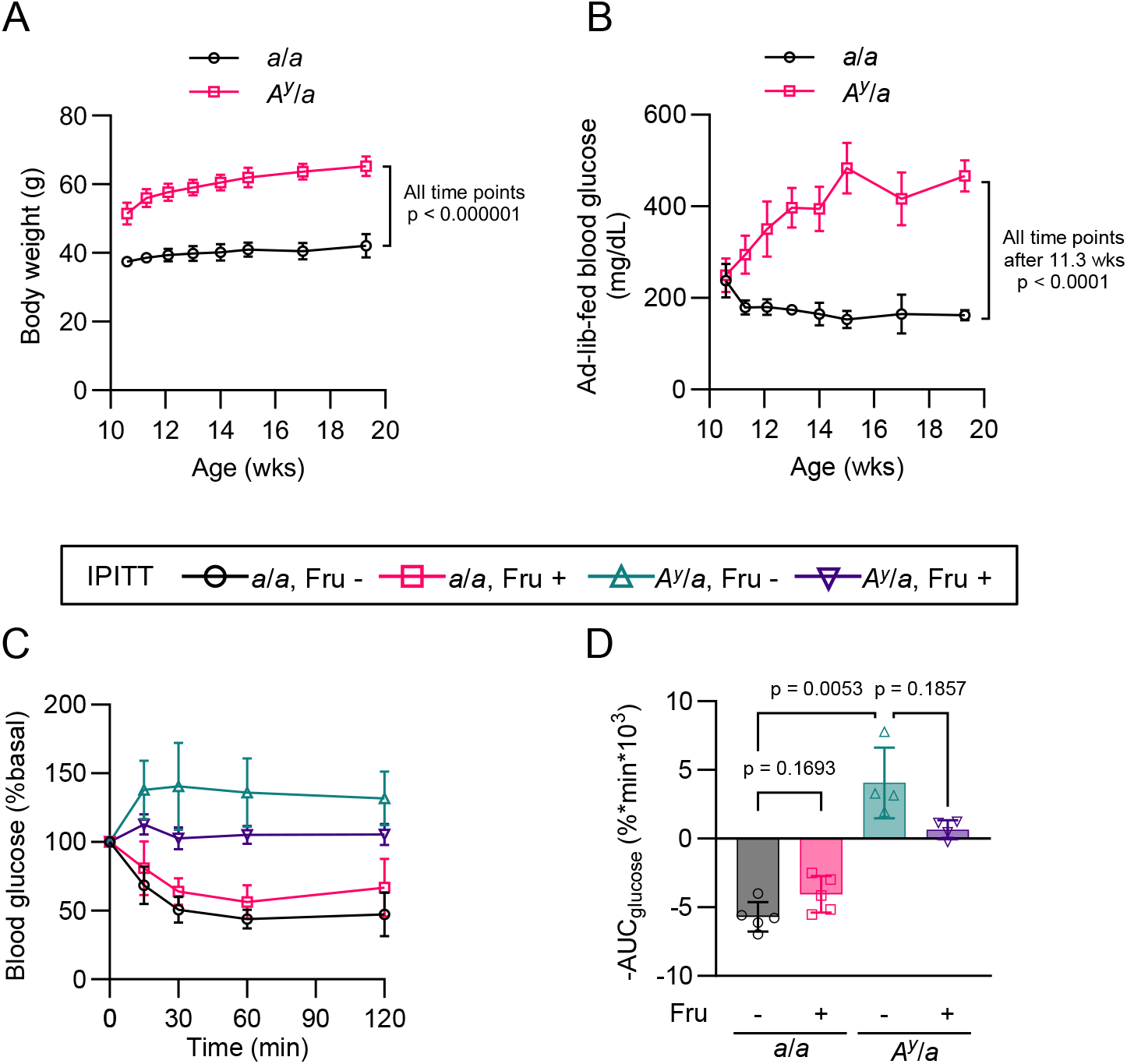
A. –B. Age-dependent changes in (A) body weight and (B) ad-lib fed blood glucose in the NSY.B6 strains. Symbols: *a*/*a*, circles; *A*^*y*^/*a*, squares. n = 6 for each strain. N = 2. C. –D. Intraperitoneal insulin tolerance test (IPITT) following 24-hour fructose ingestion. C, Blood glucose levels are expressed as a percentage at 0 min. D, AUC of blood glucose levels. *a*/*a* Fru – (circles): n = 5; *a*/*a* Fru + (squares): n = 5; *A*^*y*^/*a* Fru – (upward triangles): n = 4; *A*^*y*^/*a* Fru + (downward triangles): n = 4. N = 2. Statistical comparisons were made using Welch’s unpaired two-tailed t-test at each age in A and B, and Welch’s one-way ANOVA with Dunnett’s post-hoc test for D.

**Figure A2.**
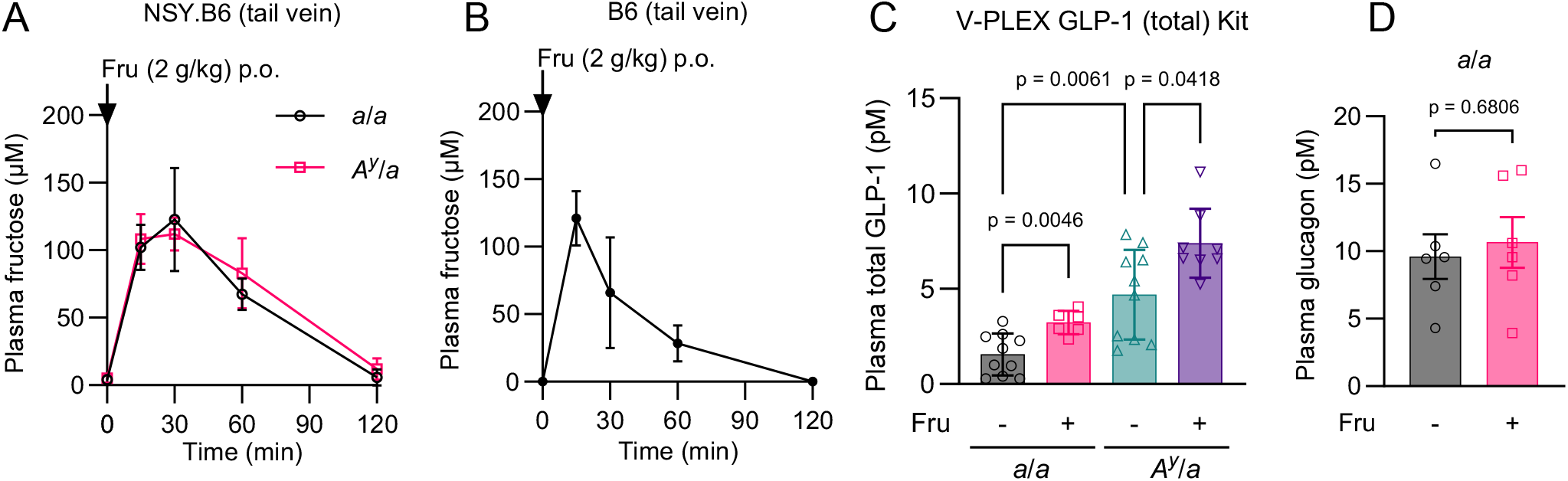
A. –B. Plasma fructose levels following oral administration of fructose (2 g/kg). (A) NSY.B6 mouse strain. n = 5 for each strain. Symbols: *a*/*a*, circles; *A*^*y*^/*a*, squares. (B) B6 mice. n = 5. N = 2. C. Plasma concentration of total GLP-1 following 24-hour fructose ingestion. Measured using Meso Scale Discovery V-PLEX GLP-1 (total) assay. *a*/*a* Fru −: n = 10; *a*/*a* Fru +: n = 6; *A*^*y*^/*a* Fru −: n = 10; *A*^*y*^/*a* Fru +: n = 8. N = 2. D. Plasma glucagon levels following 24-hour fructose ingestion in *a*/*a* mice. n = 5. N = 2. Data are presented as mean ± SD. Statistical comparisons were made using Welch’s one-way ANOVA with Dunnett’s post-hoc test for C, and Welch’s unpaired two-tailed t-test for D.

**Figure A3.**
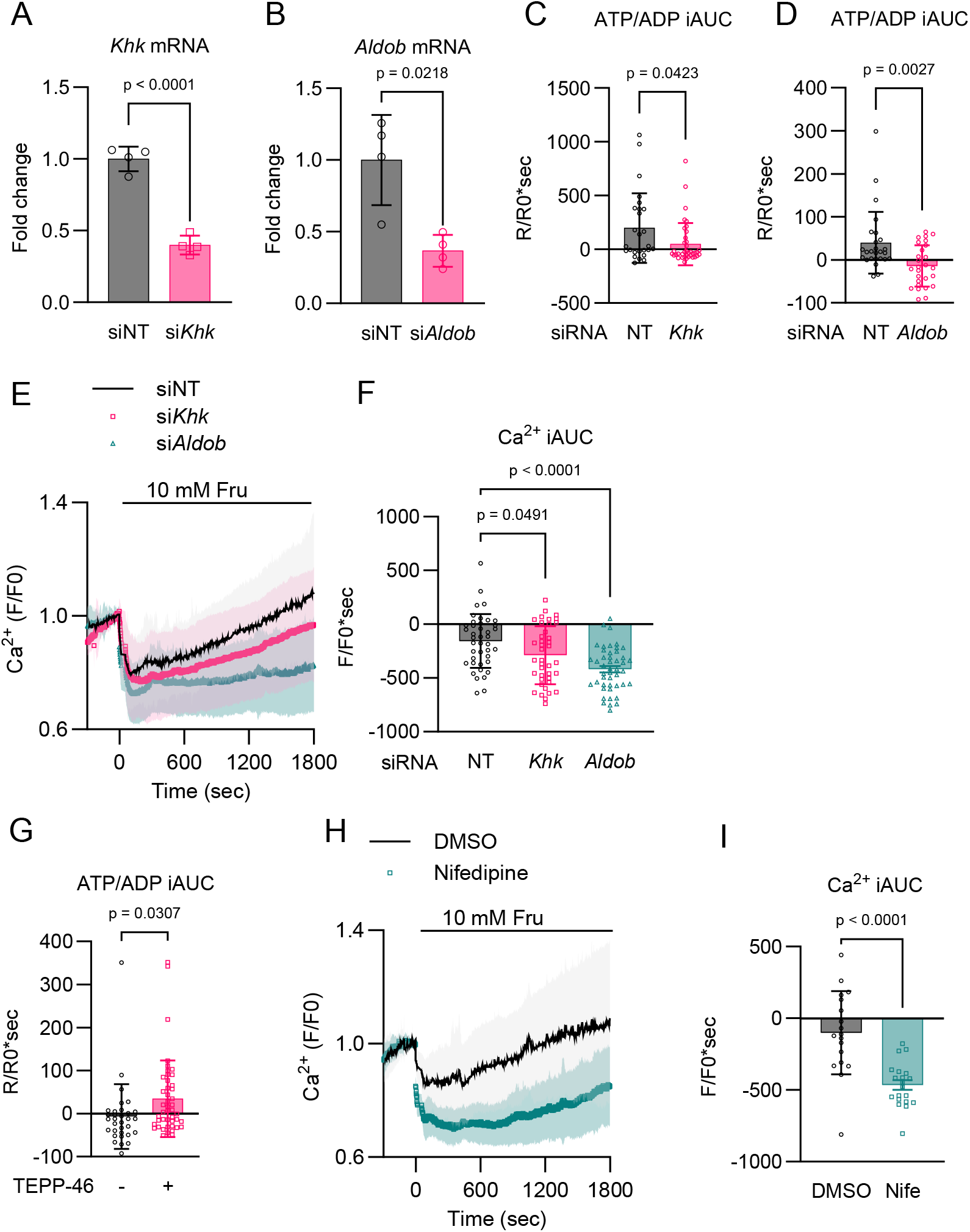
A. –B. Knockdown efficiency of (A) *Khk* and (B) *Aldob* was assessed by RT-qPCR. mRNA levels were normalised to those in siNT (non-targeting siRNA)-treated cells. n = 4, N = 2. C. –D. Effect of *Khk* and *Aldob* knockdown on fructose-stimulated ATP/ADP ratio in GLUTag cells. iAUC of Perceval-HR ratiometric signals (R/R0) are indicated in B (NT: n = 26; *Khk*: n = 38) and C (NT: n = 26; *Aldob*: n = 27), respectively. N = 2. The original values are shown in Figs. 4B and 4C. E. –F. Effect of *Khk* and *Aldob* knockdown on fructose-stimulated [Ca^2+^]i changes was visualised using Fluo-4. E, Time course of the averages of the normalised fluorescence intensity (F/F0) is indicated. For representative traces, see Fig. 4D. F, the magnitude of [Ca^2+^]i responses was quantified as iAUC using F/F0 =1 as the baseline. siNT (solid lines): n = 41; si*Khk* (squares): n = 40; si*Aldob* (triangles): n = 46. N = 2. G. Effect of TEPP-46 on the ATP/ADP ratio in GLUTag cells. iAUC of Perceval-HR ratiometric signals (R/R0) are indicated. For the original values, see Fig. 4F. TEPP-46 −: n = 32; TEPP-46 +: n = 45. N = 2. H. –I. Effect of nifedipine (Nife) on fructose-stimulated intracellular Ca^2+^ changes was visualised using Fluo-4. H, Time course of the averages of the normalised fluorescence intensity (F/F0) is indicated. For representative traces, see Fig. 4D. I: The magnitude of [Ca^2+^]i responses was quantified as iAUC using F/F0 =1 as the baseline. DMSO (solid lines): n = 18; Nife (squares): n = 20. N = 2. Data are presented as mean ± SD. Statistical comparisons were made using Welch’s one-way ANOVA with Dunnett’s post-hoc test for F, and Welch’s unpaired two-tailed t-test for the other results.

**Figure A4.**
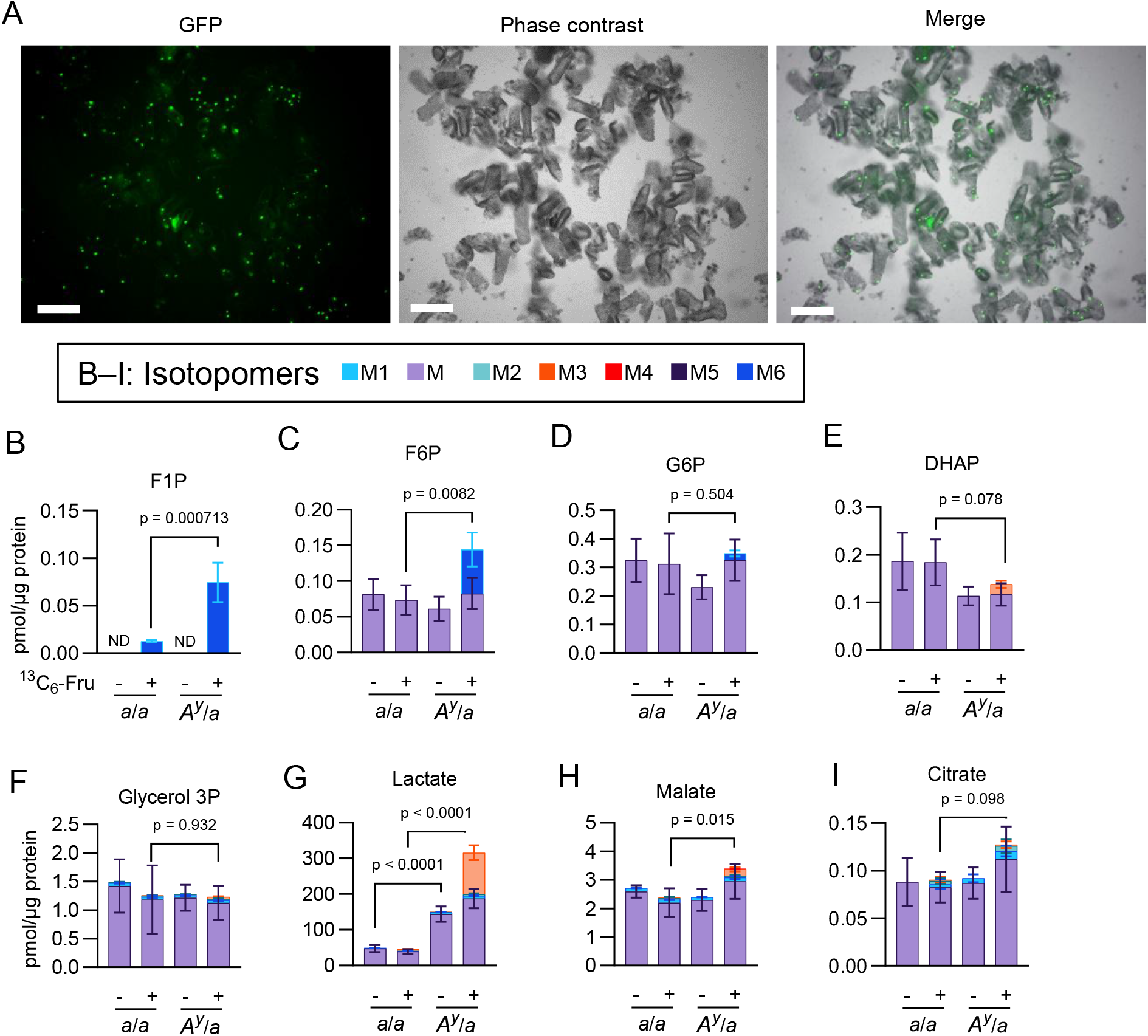
A. Fluorescence microscopy images of intestinal crypts freshly isolated from *Gcg*^gfp/+^ mice. Scale bars, 1mm. B. –I. ^13^C_6_-fructose tracing in intestinal crypts freshly isolated from *a*/*a* and *A*^*y*^/*a* mice. n = 4. The total content and isotopic distribution of each intermediate are presented as stacked bar graphs. n = 4. N = 2 Statistical comparisons were made between the sum of all indicated isotopomers using Welch’s unpaired two-tailed t-test.

